# Extrachromosomal DNA Associates with Nuclear Condensates and Reorganizes Chromatin Structures to Enhance Oncogenic Transcription

**DOI:** 10.1101/2024.09.17.613488

**Authors:** Aziz Taghbalout, Chia-Hao Tung, Patricia A. Clow, Ping Wang, Harianto Tjong, Chee Hong Wong, Diane D. Mao, Rahul Maurya, Meng-Fan Huang, Chew Yee Ngan, Albert H. Kim, Chia-Lin Wei

## Abstract

Extrachromosomal, circular DNA (ecDNA) is a prevalent oncogenic alteration in cancer genomes, often associated with aggressive tumor behavior and poor patient outcome. While previous studies proposed a chromatin-based mobile enhancer model for ecDNA-driven oncogenesis, its precise mechanism and impact remains unclear across diverse cancer types. Our study, utilizing advanced multi-omics profiling, epigenetic editing, and imaging approaches in three cancer models, reveals that ecDNA hubs are an integrated part of nuclear condensates and exhibit cancer-type specific chromatin connectivity. Epigenetic silencing of the ecDNA-specific regulatory modules or chemically disrupting liquid-liquid phase separation breaks down ecDNA hubs, displaces MED1 co-activator binding, inhibits oncogenic transcription, and promotes cell death. These findings substantiate the *trans*-activator function of ecDNA and underscore a structural mechanism driving oncogenesis. This refined understanding expands our views of oncogene regulation and opens potential avenues for novel therapeutic strategies in cancer treatment.

## Introduction

Extrachromosomal circular DNA (ecDNA) has emerged as a critical oncogenic alteration in cancer genomes with significant implications for patient outcomes and tumor progression^1,2^. EcDNA is widely prevalent across various cancer types and its prevalence increases dramatically from diagnosis to metastasis, reaching up to 48% in metastatic cancers^3,4^. Furthermore, ecDNA occurrence is predominantly associated with the most aggressive cancer types, including glioblastoma (GBM) and sarcoma^1^, and strongly linked to poorer patient prognoses^5^. Hence, ecDNA can be considered a hallmark of advanced cancer^1^. As a *bona fide* mechanism for oncogene amplification^4,6^, ecDNA contributes to cancer progression by offering cancer cell growth fitness and promoting intratumoral heterogeneity (ITH)^3,7-10^. The oncogenes most commonly found on ecDNAs include *CDK4*, *EGFR*, *ERBB2*, *MDM2*, and *MYC*^1^.

EcDNAs exist as chromatinized DNA circles within the nucleus, display a highly accessible chromatin structure, and tend to cluster together to form transcription hubs with activator complexes, which facilitate increased transcriptional activity^11,12^. Such unique chromatin conformation provides a topological framework critical for its function in cancer^11,13-16^. Our previous study showed that ecDNA makes extensive chromatin interactions with chromosomal genes in patient tumor-derived ecDNA (+) GBM cells^15^. In deciphering the ecDNA–chromosome interactomes, we observed the interaction sites exhibiting the key characteristics of super-enhancers (SEs), large clusters of enhancers densely bound by transcription factors and coactivators, that drive high levels of transcription in tumor cells^15^. Moreover, the genes connected to ecDNAs display increased transcriptional activity, and many of these genes are known oncogenes. Collectively, these observations lead to a conceptually exciting model that ecDNAs act as potent and versatile mobile trans-activators and elicit target-specific chromatin contacts in *trans* to regulate global gene expression. These unique activator activities of ecDNA, coupled with its random segregation during cell division^11^, have potential functional implications in promoting cell proliferation, driving ITH and tumor evolution^2,13,15^.

While the concept of extrachromosomal enhancers has garnered significant attention, this model is primarily based on the observed correlation between enhancers on ecDNAs and the elevated transcription activities of their associated chromosomal genes from analyses largely confined to GBM cancer models. It is imperative to directly establish the causal links between ecDNA enhancers and increased gene expression across a broader range of cancer types, validate the specificity of the ecDNA-mediated chromosomal interactions, and substantiate the significance of this mobile enhancer function in oncogenic proliferation. In this comprehensive investigation, we employed a diverse array of advanced genomic approaches including multi-omics profiling, epigenetic editing, chemically induced perturbation, and live-cell imaging, to elucidate the precise mechanisms underlying ecDNA mobile enhancer activity and its specificity and impact on chromatin organization. Our data provide compelling and direct evidence that substantiates the broader applicability of the ecDNA enhancer model across diverse biological contexts. Notably, we reveal that ecDNA-aggregated hubs are part of phase-separated nuclear condensates formed through liquid-liquid phase separation (LLPS), a process known to compartmentalize nuclear activity^17,18^. By disrupting the ecDNAs’ chromatin conformation and perturbing their enhancers, we unambiguously demonstrate the effects on targeted gene transcription and cellular proliferation. This study offers robust support for a chromatin conformation-based transcriptional mechanism, profoundly expands current views of how oncogenes are regulated epigenetically to impact tumor behavior and reveals potential therapeutic interventions targeting ecDNA enhancers in cancer.

## Results

### Mapping multiplexed ecDNA-chromosomal interactions at single molecule resolution

EcDNA has been proposed to promote oncogenic transformation through an unconventional mobile enhancer model^15^ in which ecDNAs are making specific contacts with chromosomal gene promoters in the nucleus to enhance their transcription. To understand the stoichiometry and complexity of the ecDNA target repertoire and their specificity across different cancer models, we harnessed the single-molecule ChIA-Drop (Chromatin Interaction Analysis by Droplet sequencing) analysis^15,19^ to systematically map ecDNA-chromosome interactions at single-molecule resolution in ecDNA-positive cancer cell lines. Different from many of the chromatin interaction assays, which relay on proximity ligation to capture pairwise contacts from bulk chromatin complexes^20,21^, ChIA-Drop leverages microfluidics to partition individual chromatin complexes for droplet-specific barcoding of multiplex chromatin fragments in contacts (Figure 1A). These fragments were pooled for sequencing, and chromosomal loci associated in each chromatin complex were grouped based on their shared barcodes to decipher multiplex chromatin contacts. EcDNA-originated chromatin interactions, and their cognate chromosomal targets can be deciphered *in vivo* at genome-wide scale.

**Fig. 1.**
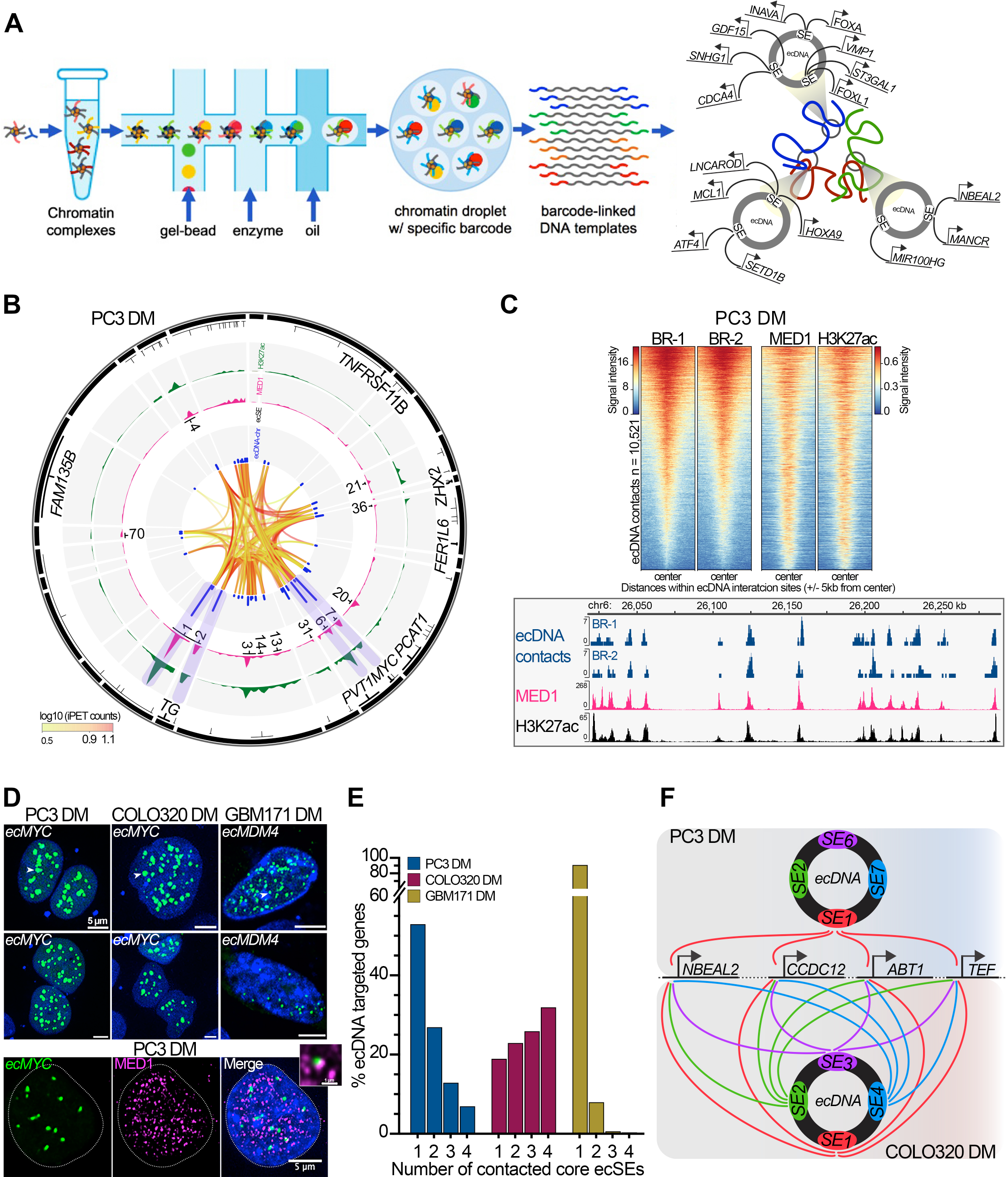
EcDNA localizes as part of MED1 nuclear condensates and forms multiplexed chromosomal interactions. **A.** Schematics of ChIA-Drop analysis. ChIP-enriched chromatin complexes are loaded onto the device to produce GEM droplets, each with a unique barcode. Barcoded amplicons are pooled for sequencing and mapping to reveal chromatin interactions associated with individual ecDNAs. **B-C.** The concordance of aggregated contact frequencies on ecDNAs and chromosomes with MED1 binding and H3K27ac intensities in PC3 DM cells. From inner to outer in the circos plot of *ecMYC* (in 10-kb resolution): interaction frequencies (IFs) of *cis*- and *trans*-loops (blue), ranked ecSEs. Fold enrichment of MED1-binding (magenta) and H3K27ac (green), genes (black). Regions of concordance are highlighted in purple. Color scale: log-transformed iPET counts. Heatmaps of the signal densities within ± 5 kb of the peaks of chromosomal sites contacting ecDNAs, MED1-binding and H3K27ac from two biological replicates (BR). Scales and numbers (n) of ecDNA-associated chromosomal contacts are shown. IGV screenshot in hg38 chr6:26,015,000-26,290,000 is shown. BR: biological replicates. **D.** Casilio live-cell imaging of PC3 DM, COLO320 DM and GBM171 DM cells shows green fluorescently labeled ec*MYC* and ec*MDM4* foci (arrowheads) within Hoechst-stained nuclei (blue) on top and co-immunofluorescence staining using MED1 antibody (magenta) on bottom. A zoomed-in view on top right highlights overlaps of MED1 droplets and *ecMYC* puncta. Scale bar as shown. **E.** The proportion of ecDNA targeted genes connected to different numbers of ecSEs with the highest chromosomal connectivity in each of the cancer models. **F.** A depiction of distinct chromosomal connectivity patterns from PC3 DM (Upper) and COLO320 DM (Lower). EcDNA-targeted genes whose promoters connected to the specified ecSEs are shown.

To capture ecDNA-chromosomal contacts associated with transcriptional activities, we produced RNA polymerase II (RNAPII)-associated ChIA-Drop datasets in three distinct cancer models, namely prostate cancer (PC3 DM)^13^, colorectal cancer (COLO320 DM)^13^, and glioblastoma (GBM171 DM). Their extrachromosomal amplification status was demonstrated previously^13,22^ or confirmed in this study by whole genome sequencing (WGS) followed by fluorescence *in situ* hybridization (FISH) (Figure S1A). These cells harbor the most frequently amplified oncogenes including *ecMYC* (PC3 DM and COLO320 DM), ec*MDM4* (<u>m</u>ouse <u>d</u>ouble <u>m</u>inute <u>4</u>), and ec*EGFR* (<u>e</u>pidermal <u>g</u>rowth <u>f</u>actor <u>r</u>eceptor) (GBM171 DM). We analyzed over 300 million paired-end sequences from each replicate and observed high consistency of the *trans-* (ecDNA-chromosomes) (Figure S1B) and *cis*-(between ecDNAs) (Figure S1C) interactions, corroborating the quality of the interactions detected. In total, we identified 14,810; 25,876; and 3,775 ecDNA-chromosomal contacts from 243,631; 573,747; and 63,101 ecDNA-associated chromatin complexes in PC3 DM, COLO320 DM, and GBM171 DM cells, respectively (Supplementary Table 1, tab 1). The highly variable number of ecDNA-associated complexes resulted from the relative ecDNA abundance in their respective cell lines, ranging from an average of 18-196 copies in reference to a diploid genome (Supplementary Table 2, tab 1). Because the sequences of ecDNAs and their native chromosomal loci are highly similar, it is extremely difficult to unambiguously distinguish the interactions mediated from ecDNAs vs. the chromosomal loci from which they originated; therefore, we adopted an orthologous Casilio-imaging^23^ approach to confirm the ecDNA-chromosomal interactions observed in the ChIA-Drop analyses. Casilio-imaging is a CRISPR-based approach that combines dCas9 and fluorescently labeled Pumilio/FBF (PUF)-tethered effectors. Using modified single guide RNAs (sgRNAs) to recruit different engineered PUF-fluorescent proteins, Casilio-imaging can simultaneously label different DNA loci in multi-color to visualize *in vivo* interactions at high resolution in individual live cells (Figure S2A). Targeted loci identified by ChIA-Drop were found to locate significantly closer to ecDNAs than randomly selected non-contacting regions from the same chromosomes surveyed from 30-40 nuclei per locus (Wilcoxon rank sum test *p*-value < 0.05) (Figure S2B), confirming the validity of detected ecDNA-chromosomal contacts.

From a total of 17,617 autosomal ecDNA-targeted genes identified by ChIA-Drop (Supplementary Table 1, tab 2), the majority (11,050 or 63%) were found in more than one cancer type, and they are enriched in cancer pathways and known oncogenes. Among the 3,447 known oncogenes collectively defined in COSMIC^24^ and the Network of Cancer Genes (NCG)^25^, 2,434 (71%) were found to be ecDNA-interacting targets including *TP53*, *ATF1,* and *MDM2* (Supplementary Table 1, tab 3). There was a core set of 2,299 chromosomal genes (CORE) tethered by ecDNAs commonly found in all three cancer lines, despite their different cancer origins and oncogene amplicons (Figure S2C). Notably, oncogenes were preferentially found (1.6-fold) in CORE compared with those uniquely identified in each of the three cancer lines. Beyond oncogenes, genes function in chromatin remodeling (1.7-fold, FDR=1.2 × 10^-7^) and chromatin organization (1.6-fold, FDR=5.6 × 10^-7^) are preferentially enriched in the CORE set as well (Supplementary Table 1, tab 4).

### EcDNA is part of nuclear condensates containing MED1 transcription coactivator

The ecDNA interactome maps display a discrete pattern where the majority of ecDNA-elicited chromatin contacts converged on a few distinct loci (Figures 1B, S1C). We further examined these distinct loci for their enhancer characteristics by performing ChIP-seq analyses to profile the H3K27ac epigenetic signatures and the binding of Mediator complex (MED1), a transcriptional coactivator. To correct for signal bias resulting from copy number amplification, we applied a unique molecular identifiers (UMIs) strategy with the input normalization^26^. In total, 38K-73K high-quality binding peaks (FDR < 0.05) of H3K27ac and MED1 were defined in each of the three cancer lines (Supplementary Table 1, tab 1), and they showed high concordance with the ecDNA interacting chromosomal loci (Figures 1C, S2D). Moreover, their aggregated contact frequencies were highly correlated with the MED1 signal intensities (Figure 1C), confirming the enhancer features of these connecting regions. As a subunit of Mediator, MED1 contains intrinsically disordered regions (IDRs) which are capable of occupying super-enhancers (SEs), known as clusters of multiple adjacent strong enhancers, to form phase-separated condensates by the liquid-liquid phase separation (LLPS) process^27^. MED1 associated condensates can compartmentalize transcription complexes and drive high-level transcription of oncogenes in many tumor cells. To visualize the conformation of ecDNA interactomes and their relative localization to MED1 in live cells, we further leveraged Casilio-imaging and MED1 co-immunostaining. In all three cancer lines, ecDNA hubs displayed discrete liquid droplets of variable sizes within the nuclei and, when co-imaged with MED1 protein, these ecDNA puncta shared significant overlap (Wilcoxon rank sum test *p*-values 1.5 × 10^-11^ and 2.2 × 10^-7^) with MED1 protein compared to random distribution (Figures 1D, S2E, Videos S1, S2), confirming that ecDNA hubs exhibit LLPS properties and are part of MED1-associated nuclear condensates. To identify potential transcription factors (TFs) involved in recruiting MED1 to ecDNA chromatin complexes, we searched for TF binding motifs specifically enriched in the MED1 binding sites within the ecDNA-chromosomal contact regions. HOMER motif analysis^28^ revealed significant enrichment of ELK1 and NRF1 binding motifs (fold of enrichment = 2.3 and 4.7, *p*-value 0 and 1 × 10^-53^, respectively), suggesting both ELK1 and NRF1 are potentially involved in recruiting MED1 to ecDNA chromatin complexes in cancer cells. Notably, ELK1 and NRF1 are key regulators of growth signaling in prostate cancer cells, and their upregulation has been associated with malignancies of prostate and colorectal cancers^29-31^.

Using the copy-number normalized MED1 binding signals, we further examined the global SE distribution by ROSE (the Ranking of Super-Enhancers) algorithm^32^. In total, 13, 23, and 24 SEs were identified on the ecDNAs (ecSEs) (Supplementary Table 2, tab 2) in PC3 DM, COLO320 DM, and GBM171 DM cells, respectively. Overall, these ecSEs are top ranked among all SEs and exhibit an order of magnitude higher MED1 binding than typical chromosomal enhancers. Among them, only a subset of ecSEs act as foci where the vast majority of the ecDNA-elicited chromatin contacts converge, suggesting the highly selective nature of ecDNA contacts (Figures 1C, S2D). To elucidate the ecDNA targeting specificity, we cross-compared the ecSE usages and ecDNA-contacting genes in different cancer models. In COLO320 DM cells, 94% of the 15,994 ecDNA-targeted genes were connected through the top four ranked ecSEs and their ecSE interaction partners are nonexclusive in nature, i.e., for a majority (> 80%), each gene was connected to multiple ecSEs (Figure 1E). Whereas in PC3 DM and GBM 171 DM cells, the majority (53% and 91%) of ecDNA-contacted genes were exclusively contacted to one ecSE (Figure 1E) (Supplementary Table 1, tab 2). Taking *NBEAL2*, *CCDC12*, *ABT1*, and *TEF* commonly found in ecSE-hubs from all ecDNA (+) cells as examples, they have individual contacts with ecSE1 in PC3 DM while concurrently connect to all four top-ranked ecSEs in COLO320 DM (Figure 1F). Such distinct stoichiometry of ecDNA-chromosomal connectivity in different cancer models implicates diverse regulatory modes for their co-activation. In PC3 DM and GBM 171 DM, a single ecSE can coordinately modulate the expression of multiple genes, whereas in COLO320 DM, multiple ecSEs function synergistically or redundantly for gene activation. Taken together, the extensive MED1 binding on ecSEs mediates the high order ecDNA-chromosomal interactions. Such MED1-based phase-separated nuclear condensates integrate ecDNA regulatory hubs with pathway-specific transcription factors and RNA polymerases to compartmentalize and coordinate gene activation.

### Rapid disruption of ecDNA hubs by agents perturbing LLPS condensates

Beyond their morphological similarity to the liquid-like droplets, ecDNA condensates are also sensitive to LLPS-interfering agents. We visualized ecDNA droplets under live-cell fluorescent microscopy by Casilio-imaging and observed that ecDNA puncta signals diminished in response to 1,6-hexanediol (1,6-HD), an aliphatic alcohol known to inhibit weak hydrophobic interactions and dissolve protein condensates^33^ in all three cancer types carrying different cargo genes (Figure 2A). As a control, fluorescent signals from the chromosomal loci where the ecDNAs originated were not affected (Figures 2A, S3A). It is worth noting that the collapse of ecDNA condensates in response to 1,6-HD displayed distinct kinetics in different cancer models. Upon exposure of 1,6-HD, the fluorescent signals from ecDNA puncta quickly faded and became weakly detectable to totally undetectable within one hour in PC3 DM and GBM171 DM cells (Figure 2A, Video S3), but only started to diminish after one hour of 1,6-HD treatment in COLO320 DM cells with some puncta signals remained visible after 155 min (Figure 2A, Video S4). Such differences in the kinetics of ecDNA condensates dissociation could reflect distinct patterns of ecDNA-chromosomal connectivity in their corresponding cell types. To verify that the loss of ecDNA fluorescent puncta was caused by the collapse of LLPS not from the reduction of ecDNA abundance, we performed real-time quantitative PCR to confirm the consistency of ecDNA copy numbers in the 1,6-HD treated cells (Figure S3B).

**Fig. 2.**
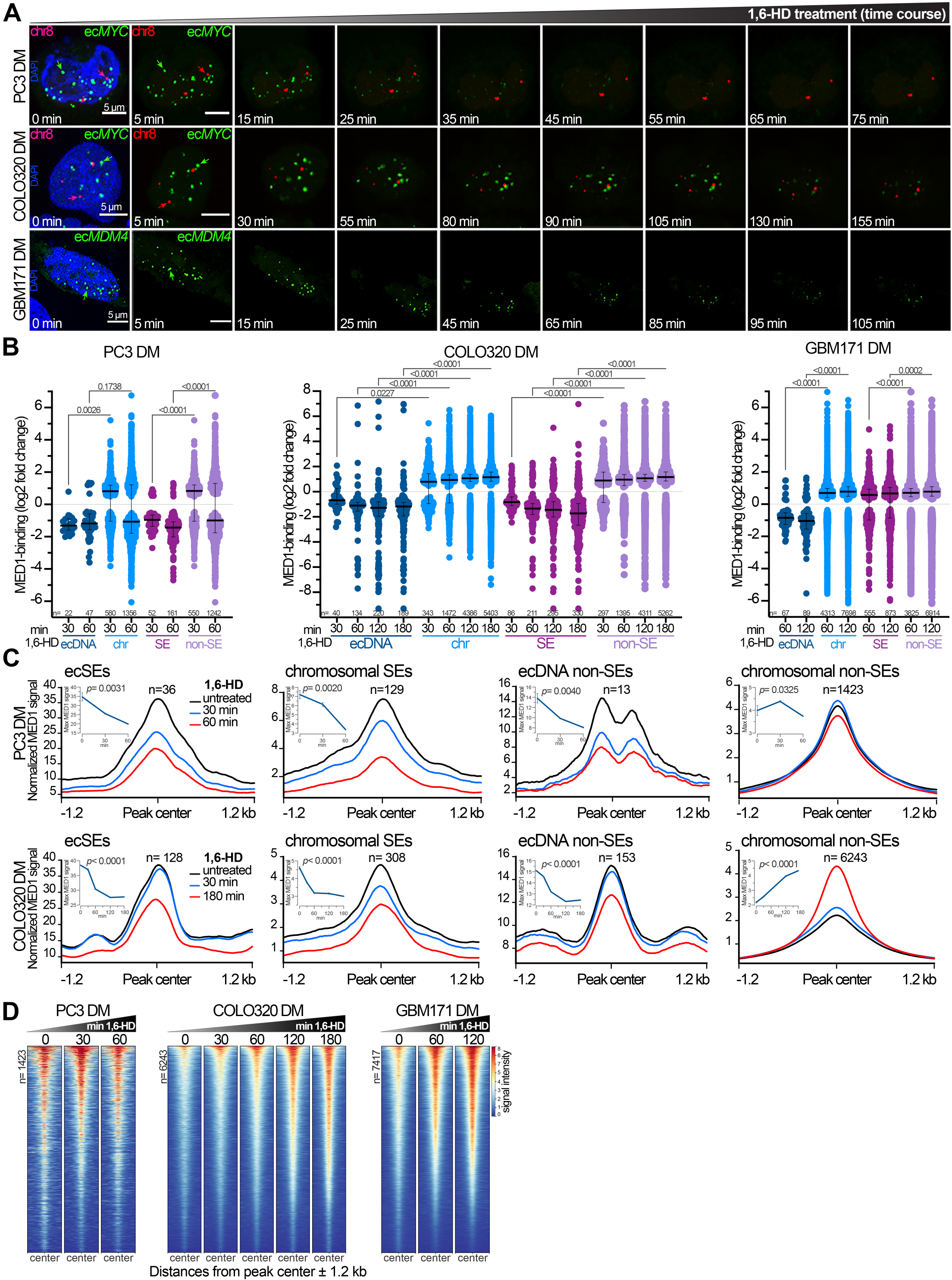
Disruption of ecDNA nuclear condensates by 1,6-HD. **A.** Casilio-imaging of the designated ecDNAs (green) in their corresponding cell lines treated with 3% 1,6-HD. Control: a chromosomal locus on chr8 (red) whose genomic coordinates are listed in supplementary Table 3. Micrographs imaged at the indicated timepoints after addition of 1,6-HD are shown. Hoechst-stained nuclei (blue) and scale bars are shown. **B.** Effects of 1,6-HD on MED1 binding. The foldchange of differential MED1 binding peaks (n) at the indicated timepoints from ecDNA, chromosomal, SE and non-SE regions are plotted with the *p*-value (one-way ANOVA) and median with interquartile **C.** Average profile plots of MED1 union peaks (n) within ±1.2 kb of the centers on SEs and outside of SEs from ecDNAs and chromosomal regions, respectively at different timepoints of 1,6-HD treatment as indicated. Maxima MED1 signal plotted over the course of 1,6-HD treatment with *p*-value (one-way ANOVA) (insert). **D.** Heat maps of the MED1 binding intensities within ±1.2 kb of the union peaks (n) on chromosomes outside of the SE regions in the cells treated with 1,6-HD at timepoints indicated.

Our results demonstrate that LLPS is the basis of ecDNA hub formation. As a key driver of LLPS, MED1 recruits ecDNAs into transcriptional condensates through binding at ecSEs. Hence, disruption of ecDNA condensates is expected to displace MED1 from ecSEs and alter ecDNA-mediated chromatin connectivity. MED1 binding was then examined by ChIP-seq analysis (Figure S4A-C Supplementary Table 1, tab1) at different timepoints throughout 1,6-HD treatment (0.5-, 1-, 2- and 3-hours). We found that MED1-binding intensities were significantly reduced on ecDNAs (one-way ANOVA, *p*-value < 0.01), with ecSEs exhibiting the greatest loss of MED1 occupancy (Figures 2B-C, S3C). Differential binding peaks (FDR ≤ 0.01) were found as early as 30 min post-treatment across all three cancer models with most of the MED1 occupancy preferentially lost on extrachromosomal loci. Notably, MED1 occupancy was regained on chromosomes during the collapse of nuclear condensates in a time-dependent manner, particularly at MED1 binding sites outside the annotated SEs (Figure 2C-D). This implies that MED1 was relocated from extrachromosomal SEs to weaker chromosomal enhancers upon LLPS perturbation, and the stoichiometry of ecSE-chromosomal connectivity could influence the kinetics of ecDNA puncta breakdown.

### Disruption of ecDNA condensates reorganizes ecDNA-mediated chromatin interactions and affects target gene expression

The repositioning of MED1 is expected to reconfigure ecSE-chromosomal contacts and dysregulate transcription. Differential chromatin interactions and gene expression were measured by *in situ* chromatin interaction (ChIA-PET)^34^ and RNA-seq analyses (Supplementary Table 1, tab1) in the 1,6-HD treated PC3 DM, COLO320 DM, and GBM171 DM cells at time points corresponding to the maximal reduction of MED1-binding. HiCRep^35^ and DESeq2 analyses demonstrated high reproducibility (<u>s</u>tratum-adjusted <u>c</u>orrelation <u>c</u>oefficient, SCC > 0.8, Pearson correlation coefficient > 0.9) between the biological replicates (Figure S4D-G). Using the normalized interaction frequencies (_n_IFs) of the ecDNA originated chromatin interactions, we performed differential analysis and revealed all significantly (|log2 Fold Change| > 1; FDR < 0.05) affected loops by 1,6-HD treatment, both in *cis* (among ecDNAs) and in *trans* (between ecDNA-chromosomes) (Supplementary Table 1, tab1). The differential loops exhibited preferentially lower _n_IFs (one-tailed Mann-Whitney *U* test, *p*-values < 0.05) in 1,6-HD treated cells, with those connected through ecSEs being affected the most. Taking PC3 DM cells as an example, 57 out of the 59-differential *cis*-interactions had reduced chromatin contacts and 53 of them were mediated from ecSEs (Figures 3A, S5A). Beyond ecDNAs, 1,6-HD also dysregulated chromosomal *cis*-interactions, particularly from SE loci (Figures 3A, S5A). Similarly, the _n_IFs of the *trans*-loops were also significantly reduced in the 1,6-HD treated cells (*p*-value = 5.8 × 10^-208^, one-tailed Mann-Whitney *U* test) in all three cancer types measured (Figures 3B, S5B). The effects of 1,6-HD on the loss of *cis*- and *trans*-loops originating from ecSE_1_ region, the reduction in MED1 binding, and the downregulation of ecSE_1_ targeted genes *HES1* and *CCN1* in PC3 DM cells are visualized in Figure 3C.

**Fig. 3.**
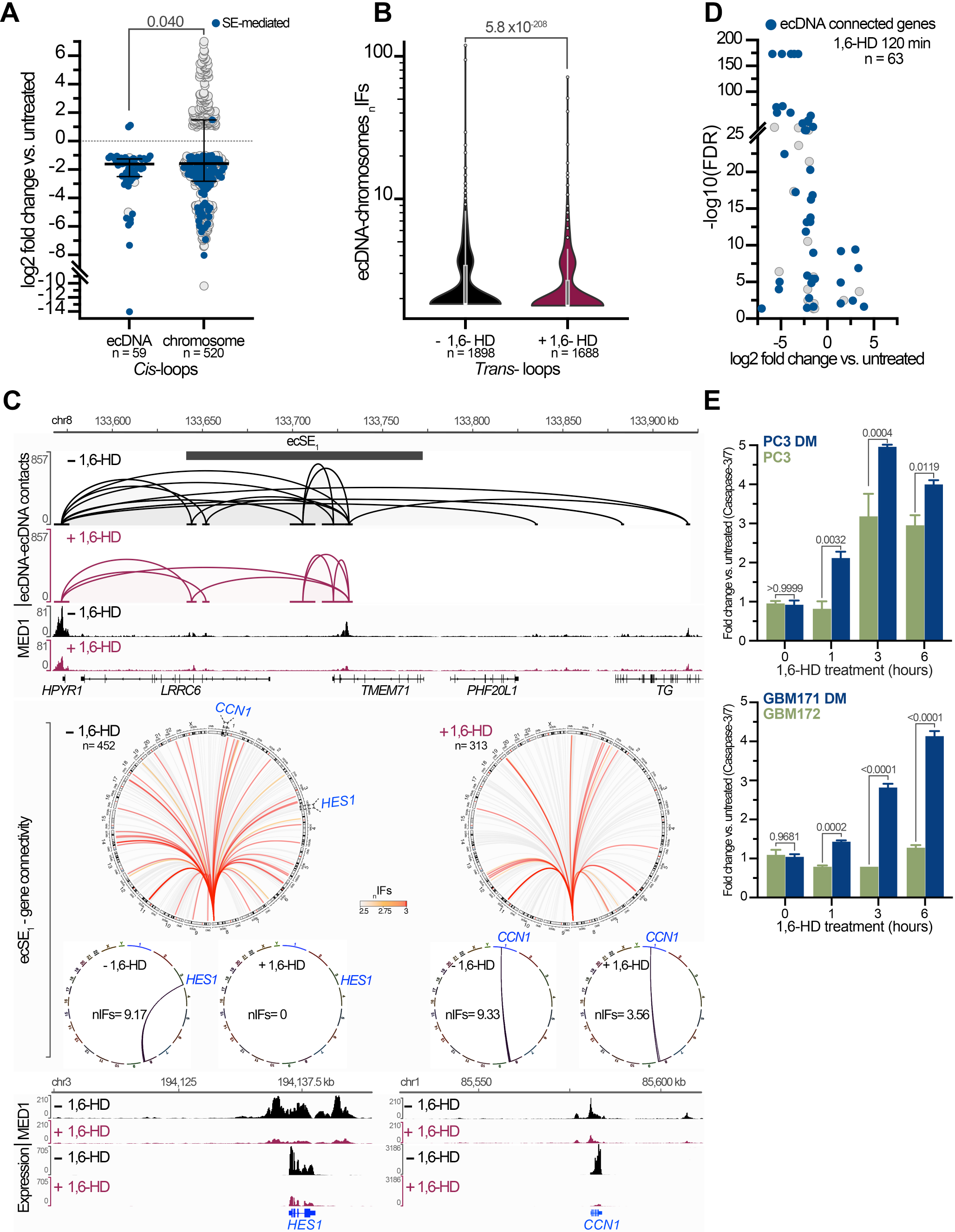
The effects of 1,6-HD on ecDNA chromatin interactions, transcription and cell viability. **A.** Log2 foldchange of the differential *cis*-loops (n) detected on ecDNAs and chromosomal loci in 1,6-HD treated PC3 DM cells. SEs are labelled in blue. *P*-values (one-tailed Mann-Whitney *U* test) and median with interquartile range are shown. **B.** _n_IFs of ecDNA-chromosomal interactions (n) in PC3 DM cells untreated (black) and treated (maroon). *P*-values (one-tailed Mann-Whitney *U* test) and median with interquartile range are shown. **C.** Screenshots of IGV browser in the region surrounding ecSE_1_ (Upper) and its chromosomal connectivity (Lower circos plots) with numbers of gene contacts (n) in PC3 DM cells. Tracks from the ecDNA-mediated chromatin contacts, MED1 binding intensities, and RNA-seq read coverages in the presence (maroon) and absence (black) of 1,6-HD are shown. Examples of two of the ecDNA target genes (*HES1* and *CCN1*) whose changes in ecDNA interaction frequency _n_IFs and RNA expression in response to 1,6-HD are shown. **D.** A volcano plot of differentially expressed genes (n) from PC3 DM treated with 1,6-HD. Genes connected with ecDNAs are shown (blue). **E.** Disruption of LLPS promotes apoptosis in ecDNA (+) cells. Fold change of Caspase 3/7 activity, as determined by flow cytometry, in ecDNA (+) cell types (blue), ecDNA (-) cells (green) and *p*-value (two-way ANOVA) at specified time points are shown.

The effects of the collapse of ecDNA condensates on transcription were evaluated by differential gene expression analyses. In the PC3 DM and GBM171 DM cells, which responded rapidly to LLPS disruption, a global downregulation of gene expression was detected within one-hour of 1,6-HD exposure. >80% of the differentially expressed genes (FDR < 0.05) were down-regulated and ecDNA-targeted genes were preferentially affected (Figures 3D, S5C-E). As examples, the _n_IFs between ecSE_1_ and the promoters of *CCN1* and *HES1*, the genes associated with promoting tumor angiogenesis^36^ and neuroblastoma metastasis^37^, were significantly reduced, and their expression were also significantly repressed (Figure 3C). In COLO320 DM, there was an initial moderate transcription activation (52 genes) at one-hour post-1,6-HD exposure, followed by progressive transcription downregulation after a prolonged four-hour exposure, mostly affecting genes connected to ecDNAs (*p*-value = 0.0028, two-way ANOVA test) (Figure S5C). This was likely due to the difference in 1,6-HD response kinetics of COLO320 DM cells and the global MED1 eviction from ecDNAs to chromosomal enhancer loci. Notably, the expression of the extrachromosomal oncogenes, like *MYC*, *PVT1*, *MDM4* and *EGFR*, was unaffected by the breakdown of ecDNA condensates, suggesting LLPS is not required to enhance ecDNA cargo gene expression.

### Disruption of LLPS promotes apoptosis in ecDNA-containing tumor cells

The aberrant chromatin contacts and transcriptional dysregulation resulting from the breakdown of ecDNA condensates ultimately affected cell viability. We quantified the percentage of apoptotic cells induced by 1,6-HD by labelling cells with a dye that specifically recognizes activated caspases 3/7 in the apoptotic pathways. There were significantly more cells undergoing apoptosis when exposed to 1,6-HD in all three ecDNA (+) cell lines (Figure S6). We leveraged the ecDNA (-) cells from corresponding prostate PC3^38^ and GBM GBM172 cancer types to assess if the elevated apoptosis rates were due to the presence of ecDNAs. The induction of apoptosis by 1,6-HD was significantly more pronounced (*p*-value < 0.001, two-way ANOVA test) in ecDNA (+) cancer cells (Figure 3E). In PC3 DM and GBM171 DM, the levels of apoptotic cells doubled within the first and third hours, respectively (*p*-value < 0.0001, two-way ANOVA test) when the general toxicity of 1,6-HD was undetectable.

Taken together, our data substantiate the ecDNA-associated chromatin condensates as the basis of how ecDNAs achieve multi-gene activation. Assembled through the ecSE-MED1 binding, ecDNA condensates can drive global transcription. Treating ecDNA (+) cells with 1,6-HD alters LLPS and destabilizes the ecDNA-associated condensates, which hampers the regulatory function of ecDNAs and promotes apoptosis. Hence, the manipulation of ecDNA-chromosomal contacts could be exploited to inhibit cancer cell proliferation.

### Casilio-i, a CRISPR-based epigenetic editing approach, is effective in silencing ecSEs

To directly establish the causal roles of ecSEs in tethering chromosomal genes, rewiring cellular transcription, and promoting tumor fitness, we deployed CRISPR-mediated epigenetic editing approaches^39^ to systematically disrupt the activities of ecSEs in the ecDNA (+) cancer models in which they were identified. Multiple CRISPR interference (CRISPRi)-based approaches, including CRISPRi^40,41^, Casilio^42^, CRISPRoff^43^, and enCRISPRi^41^ have been reported to repress enhancer signals. Given the multi-copy nature of ecSEs, their activities could be more resilient to conventional CRISPR-based assays^16^. Therefore, a robust editing assay that can deliver high efficacy in enhancer silencing is needed. We conducted a systematic evaluation of the interference efficiency between several CRISPRi-based approaches and Casilio-i, an improved version of CRISPRi^42^. Different from CRISPRi, which recruits a *single* dCas9-KRAB fusion protein to modify targeted enhancers with repressive H3K9me3 histone mark, Casilio-i leverages a programmable Pumilio RNA-binding (PUF) protein to recruit *multiple copies* of KRAB repressors to targeted loci through sgRNAs attached with multiple (5x) PUF-binding RNA sequences (PBS) (Figure S7A), hence amplifying its epigenetic silencing activities. To determine if additional epigenetic repressive factors, like LSD1^41^ or MeCP2^44^, can achieve higher and more stable epigenetic silencing, we also assessed the enhanced CRISPRi (enCRISPRi) and Casilio-i+ methods (Figure S7A). Isogenic cell lines carrying different transgenes stably expressing dCas9, dCas9-KRAB and dCas9-LSD1, along with their associated KRAB fusion proteins (Figure S7A) under doxycycline (Dox)-inducible promoters were established. The epigenetic silencing effects of Casilio-i, Casilio-i+, CRISPRi, and enCRISPRi were evaluated using a previously characterized *MYC* enhancer (hg38_chr8:127960023-127961036)^40^ by transient transfection of the validated sgRNAs. We then evaluated the reduction of H3K27ac at this locus by CUT&Tag^45^ and *MYC* expression by RT-qPCR upon Dox induction. As shown in Figure S7B, Casilio-i edited cells exhibited significantly lower (*p*-value < 0.0001, one-way ANOVA test) levels of H3K27ac and *MYC* expression in a Dox dependent manner than cells subjected to the other CRISPR interference approaches. Notably, epigenetic editing by dual repressor proteins did not yield better silencing effects, presumably due to the lower efficiency in expressing dCas9-fusion proteins of larger sizes.

We then applied Casilio-i to epigenetically perturb the four ecSEs with the highest chromosomal connectivity identified in prostate and colorectal cancer models, namely ecSE_1_, ecSE_2_, ecSE_6,_ and ecSE_7_ in PC3 DM and ecSE_1_, ecSE_2_, ecSE_3_ and ecSE_4_ in COLO320 DM (Figures 1B, S2D), respectively. Dox-inducible Casilio-i engineered PC3 DM and COLO320 DM cell lines were transiently transfected with sgRNAs (Supplementary Table 3, tab2) targeting selected ecSEs, individually or all four combined (ecSE_combo_). The effectiveness of the silencing was evaluated by quantifying H3K9me3 and H3K27ac signals across the targeted loci (Figures 4A, S7C) and the specificity was further confirmed by genome-wide analysis (Figure S7D-E). To ensure validity and reproducibility, three independent transfections were analyzed for each edited ecSE target. Overall, we detected massive depositions of repressive H3K9me3 and depletion of activating H3K27ac on the designated ecSE regions in their respective cells.

**Fig. 4.**
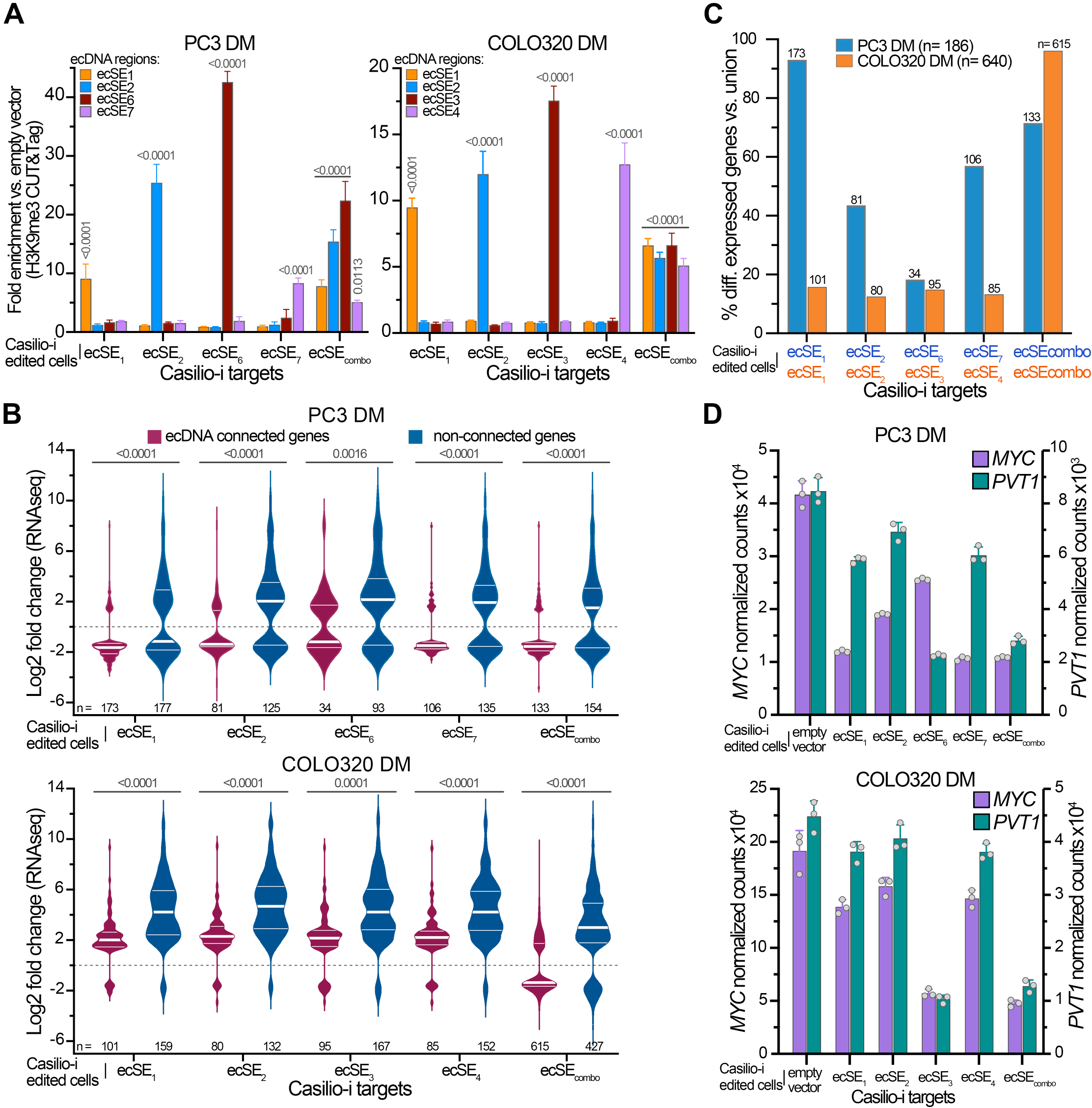
Silencing of ecSEs dysregulates global gene expression. **A.** H3K9me3 fold enrichment in specified ecSE regions, individually or in combination (ecSE_combo_) targeted by Casilio-i in PC3 DM (left) and COLO320 DM (right). *P-*values (two-way ANOVA) are shown. **B.** Violin plots of log2 foldchange of differentially expressed genes (n) (FDR < 0.05) between ecDNA targets (maroon) and non-contacted chromosomal genes (blue) in PC3 DM (Upper) and COLO320 DM (Lower) cells subjected to different ecSE silencing as indicated. Median with interquartile range and *p*-values (Mann-Whitney *U* test) are shown. **C.** Percentages of differentially expressed genes vs. union of total affected by each individual ecSE and ecSE_combo_ Casilio-i targeting in PC3 DM and COLO320 DM cells. **D.** Normalized counts of *MYC* and *PVT1* transcripts in PC3 DM (Upper) and COLO320 DM (Lower) cells subjected to silencing of specified ecSEs. Mean and standard deviation bars are shown.

### Perturbing ecSEs interferes with ecDNA-associated chromatin connectivity and transcription

To dissect the functional impacts of disrupting ecSE activities on global transcription, we evaluated the differentially expressed genes (FDR < 0.05) by RNA-seq analysis (Supplementary Table 1, tab1, Figure S4 H-I). Hundreds of dysregulated genes were detected in the PC3 DM and COLO320 DM cells in which the individual ecSEs or ecSE_combo_ activities were disrupted. In PC3 DM, chromosomal genes contacted by ecDNAs were preferentially downregulated (*p*-value < 0.0001, Mann-Whitney *U* test) (Figures 4B, S8A) and epigenetic silencing of ecSE_1_ exerted the largest effects. Out of the total 186 dysregulated ecDNA targets, 173 (93%) were found in ecSE_1_ Casilio-i cells and the effects from silencing of ecSE_2_, ecSE_6,_ and ecSE_7_ are mostly redundant (Figure 4C), indicating that ecSE_1_ is the dominant driver of ecDNA’s mobile enhancer activity. In COLO320 DM cells, when the individual ecSEs were inactivated, only a small subset (13-16%) of the totally dysregulated ecDNA targets were affected and the effects were largely upregulated (Figure 4B-C), potentially caused by shifting MED1 to other ecSE loci. Only when the enhancer activities of all four ecSEs were abolished (ecSE_combo_) the transcriptional effects were mostly significantly downregulated (*p*-value < 0.0001, Mann-Whitney *U* test), with dysregulation of 615 out of the total 640 (96%) ecDNA-targeted genes, reflecting the redundant nature of the ecSEs in modulating transcription. Such exclusive vs. synergistic activities of ecSEs observed in PC3 DM and COLO320 DM cells, respectively, are consistent with their chromosomal connectivity patterns (Figure 1E-F). Gene functions in DNA replication, regulation of chromosome organization, and ribosome biogenesis are most significantly affected (*p*-value < 7 × 10^-9^) in the ecSE-inactivated cells (Figure S8B).

The disruption of ecSEs also suppresses transcription of ecDNA cargo genes, as evident by the steep and coordinated downregulation of *MYC*, *PVT1*, *TG,* and *POU5F1B* in ecSE-silenced cells (Figures 4D, S9A), supporting their *cis*-regulation mechanism^16^. While antagonistic expression patterns have been observed for *MYC* and *PVT1* in cancers driven by *MYC* mis-regulation^46^, concomitant repression of *MYC* and *PVT1* was achieved by silencing the ecSEs in both ecDNA (+) cancer types (Figure 4D), implying that extrachromosomal genes are subject to distinct regulatory conformation from their chromosomal counterparts. It is worth noting that such a drastic reduction of *MYC* and *PVT1* transcripts (20-25%) has not been achieved through manipulations of any *cis*-regulatory elements, raising an intriguing possibility that ecSEs can be exploited as a vulnerability for therapeutic strategies.

To corroborate our working model where ecDNA chromatin contacts are the basis of its enhancer function to modulate transcription, we measured ecDNA-chromosome interactions in the Casilio-i perturbed cells. Due to the limited quantities of transiently transfected cells with sgRNA, we adopted the ChIATAC chromatin interaction assay^47^ which is specifically designed for low-cell inputs. ChIATAC uniquely combines proximity ligation and transposase tagmentation deployed in the ChIA-PET and ATAC-seq to efficiently capture interactions between regulatory elements for sequencing analysis. We analyzed Casilio-i PC3 DM cells targeting ecSE_1_ and all four ecSEs (ecSE_combo_) which demonstrated the most impact on transcription. Both *cis*-(among ecDNAs) and *trans*-loops (between ecDNAs and their chromosomal targets) were identified, and their normalized interacting frequencies (_n_IFs) were found significantly lower (*p*-value < 1 × 10^-15^, one-tailed Mann-Whitney *U* test) in response to ecSE epigenetic silencing (Figure 5A). The diminishing of 3D contacts is also reflected in the intensities of aggregated _n_IFs of the *trans*-loops elicited from ecSEs in control, ecSE_1,_ and ecSE_combo_ Casilio-i cells (Figure 5B). Examples of ecDNA-mediated *cis*- and *trans*-interactions visualized by integrated genome browsers (IGV) between controls and ecSE-edited are shown in Figure 5C-D. *DNMT3B*, an ecSE_1_/ecSE_2_ target gene encoding a *de novo* methyltransferase involved in abnormal methylation in cancer^48,49^, lost all its interactions with ecDNAs in cells when all four ecSEs were silenced and lost 60% of its expression (*p*-value = 0.017). Similarly, *CDCA5*, an ecSE_1_/ecSE_7_ targeted gene encoding Sororin, which is required for cell cycle regulation and promoting the progression of malignancies such as prostate cancer^50^, lost all of its ecSEs contacts and 72% of its expression (*p*-value = 1.93 × 10^-35^) (Figure S9B). The decline in the frequency of *cis*-loops connecting ecSE_1_ and *TG* promoter, an ecDNA cargo gene, resulting from ecSE_1_-silencing also drastically down-regulated *TG* expression by 90% (*p*-value = 1.88 × 10^-190^).

**Fig. 5.**
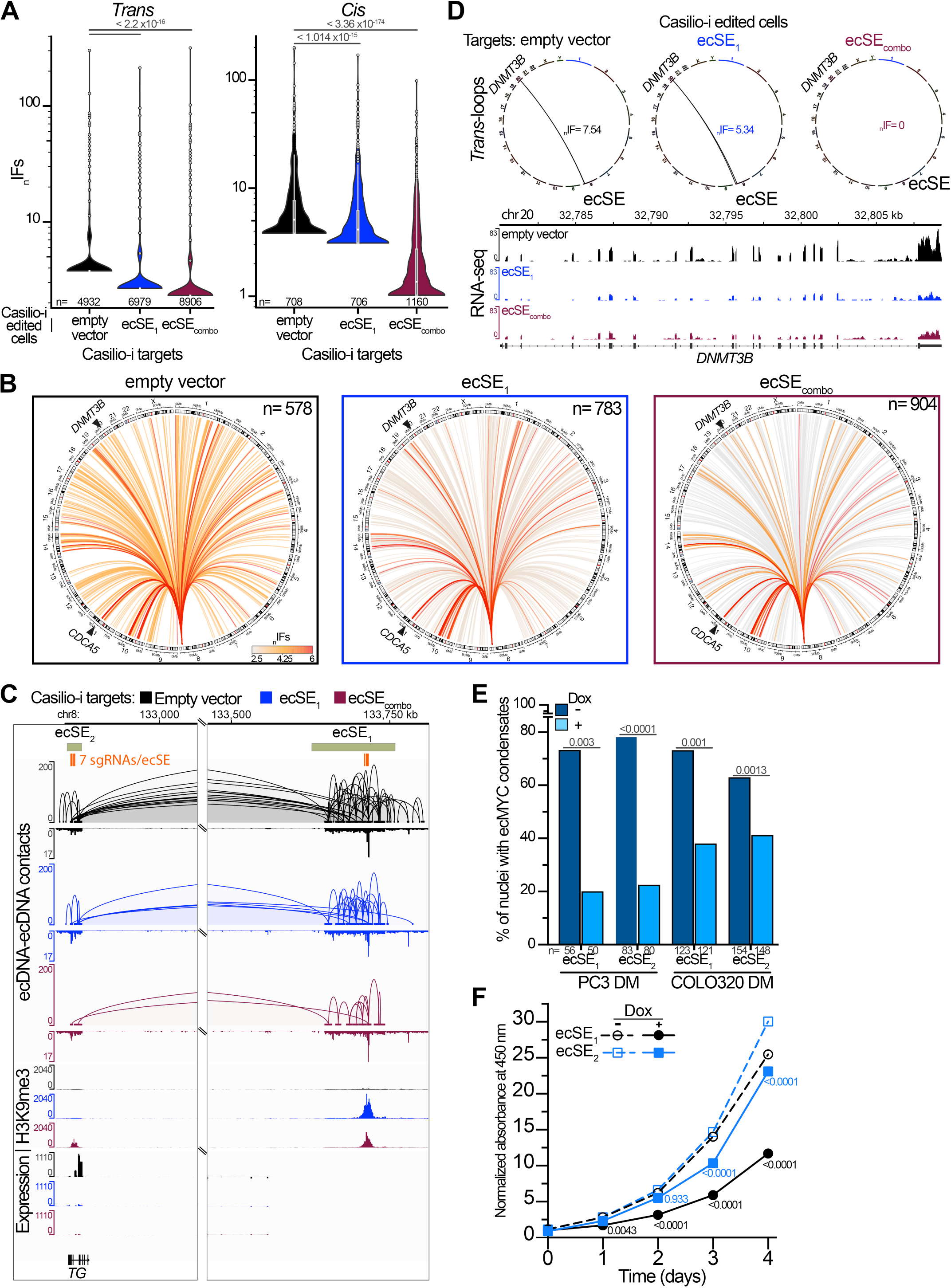
Silencing of ecSEs perturbs ecDNA-associated chromatin interactions, dissociates ecDNA condensates and inhibits cell proliferation. Control (black), Casilio-i targeted ecSE_1_ (blue), and ecSE_combo_ (maroon) in PC3 DM derived cells, **A.** Normalized interaction frequencies (_n_IFs) of ecDNA *trans-* (Left) and *cis-* (Right) loops (n). *P*-values (one-tailed Mann-Whitney *U* test) are shown. **B.** Circos plots of ecSE_1_-chromosomal interactions (n) in control, Casilio-i ecSE_1_ and ecSE_combo_ targeted PC3 DM cells. _n_IFs shown as color scale, genes exemplified in panel D and Fig. S9B are labelled. **C.** Screenshots of IGV browser from regions spanned between ecSE_1_ and ecSE_2_. Tracks (from top): *cis*-loops with ATAC peaks, H3K9me3 normalized signal intensities, RNA-seq normalized read coverage and annotated genes. Locations of ecSE-targeting sgRNAs (orange), ecSEs, and hg38 genomic coordinates are shown. **D.** Circos plots of ecSE-*DNMT3B* interactions (_n_IFs) in control, Casilio-i ecSE_1_ and ecSE_combo_ targeted PC3 DM cells. Screenshot of IGV browser of *DNMT3B* RNA-seq normalized read coverages. **E.** Percentages of nuclei containing *ecMYC* puncta in control (-Dox) and ecSE_1_ and ecSE_2_ silenced (+Dox) cells. Number of analyzed nuclei (n) and *p*-value (two-tailed Mann-Whitney *U* test) are shown. **F.** Proliferation rates (represented by normalized OD_450_) of Dox-inducible Casilio-i PC3 DM cells that stably express sgRNAs targeting ecSE_1_ or ecSE_2_ across 4-days in presence or absence of Dox, *p*-values (two-way ANOVA) are shown.

### EcSEs are required for ecDNA condensate formation and promoting tumor cell growth

The ecDNA chromatin hubs, driven by the ecSEs-MED1 association, are part of phase-separated nuclear condensates. Hence, the disruption of ecSE enhancer activities by Casilio-i is expected to breakdown ecDNA condensates. To visualize ecDNA puncta in cells with knocked-down ecSE activity, we imaged engineered Casilio-i PC3 DM and COLO320 DM live-cells that stably express the specified sgRNAs targeting ecSE_1_ or ecSE_2_. In the presence of Dox, the number of nuclei containing *ecMYC* puncta was significantly reduced (*p*-value < 0.001, two-tailed Mann-Whitney *U* test), with up to four-fold fewer nuclei displaying fluorescent ecDNA droplets (Figure 5E). The loss of ecDNA droplet signals was not due to any decay of ecDNA molecules, as ecDNA copy numbers were unaffected by Casilio-i perturbation (Figure S9C).

Consistent with the downregulation of genes involved in DNA replication (Figures S8B, S9B), cell proliferation was affected by ecSE silencing. The growth rates of the Dox-inducible Casilio-i PC3 DM cells stably expressing sgRNAs targeting ecSE_1_ or ecSE_2,_ as measured by a colorimetric-based assay^51^, were significantly lower (<50%, *p*-value ≤ 0.01, two-way ANOVA test) in the presence of Dox over the four day log-growth phase (Figure 5F), with stronger growth inhibition associated with ecSE_1_ perturbation, indicating a slow growth phenotype associated with the inhibition of ecDNA super enhancers.

Taken together, ecDNA chromatin hubs are part of phase-separated nuclear condensates. Driven by the ecSE-MED1 association, ecDNAs can act as a target-specific *trans*-activator to enhance the transcription of chromosomal genes. Both 1,6-HD and ecSE perturbation resulted in the dissociation of MED1 binding from the co-amplified enhancers and preferential down-regulation of ecDNA-targeted genes, including genes known to be involved in cell cycle and proliferation. These results directly demonstrate a structure-based regulatory function for ecDNAs in enhancing transcription and growth fitness.

## Discussion

In this study, we applied a collection of advanced chromatin profiling, genome editing, and imaging assays to advance our understanding of ecDNA function in diverse types of cancer models. We expanded the utility of ChIA-Drop to reveal multiplexed ecDNA–chromosome interactions at single molecule resolution and deeply characterized ecDNAs’ regulatory modules and target gene repertoire. We employed Casilio-imaging to enable high resolution dual-imaging of the ecDNA-MED1 aggregates and demonstrated the epigenomic editing of Casilio-i for the robust disruption of ecSE activities. The multi-omics analysis of ecSE-perturbed cells further establishes the causality between ecDNAs, their chromatin connectivity, and the regulation of chromosomal gene expression.

Intensive effort has been made to understand the mechanisms by which ecDNA modulates tumor growth, fitness, and therapeutic responses. The notion that an LLPS-based mechanism is involved in the ecDNA mobile enhancer model provides a physical basis for ecDNAs to achieve genome compartmentalization for coordinated transcriptional co-activation. LLPS is a physicochemical process shown by many to mediate the formation of numerous biological condensates^52^ and has been proposed as a mechanism promoting 3D genome organization. Here, the ecSEs act as foci where multiple genes cluster into ecDNA-associated condensates to promote local DNA contacts. Among different cancer models, distinct modes of ecSE connectivity, either highly exclusive or redundant and synergistic, were observed. Such “structural-based” ecDNA liquid condensates can be considered as cancer-specific traits, that can be exploited for therapeutic strategies. Currently, no cancer therapies directly target ecDNAs, and the ability to effectively inhibit these traits is expected to offer conceptually new opportunities for cancer therapy. Here, we demonstrated that by epigenetic modulation of a small segment of extrachromosomal regulatory elements, ecSE perturbation can disintegrate the ecDNA condensates and result in transcriptional dysregulation of specific oncogenes, even those currently considered undruggable.

It is important to note that distinguishing ecDNAs from their native chromosomal loci is exceptionally challenging due to their nearly identical sequences. This unresolved challenge highlights the pressing need for new tools and technologies to address this technical gap. As a result, the observed chromatin interactions and impacts from ecSE perturbation may represent aggregated effects from both chromosomal and extrachromosomal loci. Because loci on ecDNAs greatly exceed chromosomal counterparts in copy number, we believe that the major signals we observed are likely contributed by ecDNA molecules. Casilio-imaging coupled with Hoechst staining was adopted as an orthologous confirmatory approach. Moreover, interactions across different chromosomal loci are generally less frequent due to constraints from the chromosomal territories^53^.

This mobile enhancer model of ecDNA’s mode of action raises an intriguing possibility that the rewiring of enhancer–promoter interactions could underlie a new type of aberrant transcriptional control to promote intratumor heterogeneity (ITH) and tumor evolution. Given their variable copy number (CN) and ability to promote transcription, ecDNAs can promote the plasticity and heterogeneity of gene regulatory circuitry and confer a selective growth advantage to cancer cells. Such *trans*-activating mechanism, although less studied, has been demonstrated in both normal physiological contexts^54^ and during viral infection. For example, the expression of olfactory receptors in the sensory neurons^55,56^ and the modulation of the viral transcriptome by the hepatitis B virus (HBV) episomes in human hepatocytes^57,58^. Further understanding of extrachromosomal DNA-mediated genome organization could represent a new paradigm in how eukaryotic genes are regulated and how structure variants and epigenetics promote tumorigenesis.

## Acknowledgments

The authors thank Drs. Edison Liu and Roel Verhaak for their feedback and comments on the manuscript, Dr. Albert Cheng for his gift of Casilio-based plasmids and technical advice, Colleen Davis for her help in revising and suggestions in writing of the manuscript, and the Scientific Services cores flow cytometry, genome technology, and microscopy at the Jackson Laboratory for their support. Research reported in this manuscript was supported by grants from the US National Institutes of Health R01 GM127531, R33 CA236681, and R01 HG011253 (CLW); and P30CA034196, R01 NS128470 (AHK).

## Author contributions

Conceptualization, C.-L.W., A.T., C.-H.T., and P.W.; Methodology, A.T., C.-H.T., P.A.C., H.T., P.W., C.H.W., C.Y.N., and D.D.M.; Formal analysis, A.T., C.-H.T., P.A.C., H.T., and C.H.W.; Investigation, A.T., C.-H.T., P.A.C., P.W., C.Y.N, M.-F.H., and R.M.; Resources, C.-L.W and A.H.K.; Writing – original draft, C.-L.W., A.T., C.-H.T.; Writing – review & editing, C.-L.W., A.T., C.-H.T, P.A.C., H.T. and A.H.K.; Funding acquisition, C.-L.W., A.H.K; Supervision, C.-L.W..

## Declaration of interests

A.H.K. is a consultant for Monteris Medical and has received research grants from Stryker for a clinical outcomes study about a dural substitute, which have no direct relation to this study. The other authors declare no competing interests.

## SUPPLEMENTAL INFORMATION

### Supplemental Figures

Figures S1-S9 with figure legends.

### Supplemental Tables

Table S1. Excel file containing sequencing summaries of all omics-based analyses and annotation of ecDNA-connected genes.

Table S2. Excel file with information on defined ecDNA regions, ecDNA copy number, ecSE locations and ROSE analysis.

Table S3. Excel file listing plasmids, sgRNAs and primers used in this study.

### Supplemental Videos

Video S1. Casilio imaging of *ecMYC* in COLO320 DM cells. Related to Figure 1D.

Video S2. Casilio imaging of *ecMYC* in PC3 DM cells. Related to Figure 1D

Video S3. Breakdown of ecDNA condensates by 1,6-HD in PC3 DM cells. *ecMYC* (green), controls from chromosomal regions (red). Related to Figure 2A

Video S4. Breakdown of ecDNA condensates by 1,6-HD in COLO320 DM cells. *ecMYC* (green), controls from chromosomal regions (red). Related to Figure 2A

## STAR METHODS RESOURCE AVAILABILITY

Further information and requests for resources and reagents should be directed to and will be fulfilled by lead contact Chia-Lin Wei.

## Data and code availability

All data described in this study are being deposited in NCBI’s Gene Expression Omnibus GSE275060. All software tools used in this study are listed in the STAR Methods description.

### Cell lines

K562^59^, COLO320 DM^22^, and PC3 DM (-)^38^ were purchased from ATCC. PC3 DM^6^ was a gift from the laboratory of Dr. Roel Verhaak at Yale school of medicine. Glioblastoma neurosphere cell lines GBM171 DM and GBM172 were generated from resected brain tumor specimens. All human study research related to this study has been approved by the Institutional Review Board (IRB #201211019 and #201111001, Washington University School of Medicine). All participants donating tissue signed informed consent prior to tissue banking. Briefly, tumors were dissociated by mincing and incubation in Accutase (A6964, Sigma-Aldrich)^60,61^. Cell suspensions were passed through a 70-μm cell strainer (352350, Falcon), and red blood cells were removed (A1049201, Life Technology) according to the manufacturer’s instructions. All cells used for the omics and functional assays were subjected to extensive characterization including WGS and FISH across multiple passages to confirm the stability of ecDNAs in these cancer models.

## Method Details

### Multi-omics assays

*Whole genome sequencing*– WGS and copy number analyses were performed on PC3 DM, COLO320 DM, and GBM171 DM cells as previously described^15^. Sequencing reads were aligned to the human hg38 reference genome assembly, and the average copy numbers were computed from the deduplicated reads using R package “readDepth”^62^.

*ChIA-Drop analysis*– RNAPII ChIA-Drop experiments were performed as previously described^15,19^. For data processing into chromatin complex interaction data, the sequencing reads were aligned to the hg38 reference genome using the 10X Genomics Long Ranger pipeline (v2.1.5), and droplet-specific barcodes were identified. Aligned reads flagged with PCR duplicates were discarded. The uniquely mapped reads with length ≥ 50 bp and MAPQ ≥ 30 were extended by 500 bps from their 3’ end, and those with the same droplet-specific barcodes within 3 kb distance were merged. We defined interaction loci as anchors detected by ≥ 3 reads and overlapped with RNAPII peaks. Multiplex interactions were defined from fragments of the same barcodes mapped to different locations. Chromatin complexes containing fragments overlapping with ecDNA regions were defined as ecDNA-associated chromatin interactions. Interactions were classified as either *cis* (among ecDNAs) or *trans* (ecDNA-chromosomal), and their interaction frequencies were determined by the number of times the anchor was observed in different ecDNA chromatin complexes defined by droplet-specific barcodes. For ecDNA-mediated chromosomal interactions, we required that each interaction be *trans* in nature, which means that only one of the two anchors overlapped an ecDNA region and the other anchor is from chromosomes. EcDNA targets were defined as the chromosomal genes whose TSS located ± 2.5 kb from the interaction anchors. To perform cross-library comparison, overlapping anchors were merged as interaction nodes and their genomic features were annotated based on SEs, MED1 binding, genes, and promoters.

*ChIP-seq analysis*– MED1 and H3K27ac ChIP-seq assays were performed as previously described^63^. Briefly, cells were crosslinked and lysed, the nuclei were sonicated, and the supernatants containing the soluble chromatin were immunoprecipitated with anti-MED1 (A300-793A, Bethyl Laboratories) or anti-H3K27ac (39133, Active Motif) antibodies. The immunoprecipitated DNAs were subjected to end-repair, A-tailing, and adaptor ligation with UMI adapters. Libraries were prepared using KAPA Hyper Prep Kit (KK8505, KAPA Biosystems) as previously described^15^ and sequenced on Illumina sequencers. Unique molecular identifiers (UMIs) and input DNA were used to normalize copy number (CN)^26^. The raw reads were quality trimmed and non-redundant reads were mapped to the hg38 genome. Reads with MAPQ ≥ 30 were deduplicated and used to identify the MED1 binding peaks and H3K27ac modification with FDR < 0.05 or otherwise indicated.

*ChIA-PET assay*– The RNAPII ChIA-PET experiments were performed as previously described^63^. Briefly, dually crosslinked cells were lysed, and the extracted nuclei were subjected to *in-situ Alu*I (R0137L, NEB) digestion followed by proximity ligation in the presence of the biotinylated bridge linkers. The ligated chromatins were fragmented, immunoprecipitated by anti-RNAPII antibodies (920102, Biolegend), Tn5-mediated tagmentation, and enriched by streptavidin. Libraries were prepared and sequenced as previously described^15^. ChIA-PET data were processed with ChIA-PET Utilities as previously described^15^. Reproducibility between replicates was confirmed with the HiCRep method^35^ using the <u>s</u>tratum-adjusted <u>c</u>orrelation <u>c</u>oefficient (SCC) method. Statistical assessment of the significance for *cis*-interactions was performed using ChiaSigScaled, a scalable reimplementation of ChiaSig^64^. Interaction clusters are defined as member size ≥ 3 (iPET-3+) and FDR < 0.05. All observed interactions were ranked according to their contact frequency (iPET counts), assigned a *p*-value relative to all other possible interactions, and their statistical significance evaluated by an established statistical model previously described^65,66^. We define the significant ecDNA-chromosomal interactions as iPET-2 and above (qnorm < 1 × 10^-7^) with RNAPII binding support at both anchors. They were further filtered using the ecDNA-contact chromosomal anchors generated from the ChIA-Drop assays. For differential *cis*-loops analysis, we applied shrinkage for log2 (fold change) estimation using lfcShrink from DESeq2^67^ and normalized iPET counts using DESeq2’s built-in count normalization approach with FDR < 0.05 and |log2(fold change)| > 1. For differential ecDNA-chromosomal interaction analysis, a matrix was built from the contact frequencies (accumulated iPET counts) from all node-node interactions and normalized by dividing the matrix with respective means. EcDNA-chromosomal interactions were then selected, and their normalized contact frequencies (_n_IFs) were used for comparison between two datasets.

*CUT&Tag analysis*– In K562 cells, CUT&Tag experiments were performed using 200,000 cells, rabbit anti-H3K27ac (39133, Active Motif), and the CUT&Tag-IT assay kit (53160, Active Motif) following the manufacturer’s instructions. DNA released from cells after tagmentation was used for qPCR normalization. For PC3 DM and COLO320 DM cell lines, CUT&Tag was performed on 50,000 nuclei by using rabbit anti-H3K9me3 (39765, Active Motif) or H3K27ac and CUTANA CUT&Tag Kit (14-1101, Epicypher) as instructed in the manufacturer’s manual except that digitonin was omitted from buffers as it is not required when nuclei are used^68^. CUT&Tag libraries were subjected to Illumina sequencing, and 5 million reads were generated on average per sample. Data were processed by trimming and aligning to the human genome (hg38), and the high-quality alignments were used for MACS2 peaks calling with a cutoff *p*-value < 0.001. Differential peak analyses were performed with DESeq2 using default parameters.

*ChIATAC analysis*– ChIATAC experiments were performed as previously described^47^ using 50,000 cross-linked cells. Reads were processed to define *cis-* (among ecDNAs) and *trans-* (ecDNA-chromosomal) interactions. To perform comparisons among different datasets, overlapped anchors were merged to form nodes. iPETs from the interactions that shared the same left/right nodes were combined and normalized with average to obtain the normalized interaction frequencies (_n_IFs). *Trans*-interactions were further selected for chromosomal nodes located ± 2.5 kb of TSS of ecDNA targeted genes identified in ChIA-Drop.

*RNA-seq*– Total RNA was isolated from cells using the RNeasy Mini Kit (74106, Qiagen), and corresponding strand-specific RNA libraries were generated and sequenced on the Illumina platform. Trimmed reads were aligned to the hg38 transcriptome with hisat2 (v 2.1.0)^69^, and gene expression analyses were performed by using the reference gene set GENCODE primary assembly annotation v36. Differential gene expression analysis was performed with R package DESeq2^67^ using the transcript quantifications from HTseq for 40,330 genes, including 19,962 protein-coding genes; 16,901 long non-coding RNA (lncRNA) genes; and 3,467 genes coding for other RNAs, including pseudogenes and miscellaneous RNAs.

### Bioinformatic analyses

*Motif analysis*– MED1 peaks residing within non-overlapping chromosomal fragments found in two or more ecGEMs from two ChIA-Drop replicates were used to search for transcription factor (TF) binding motifs. The peak summits were extended from their center by ± 0.5, 1, or 2 kb and scanned for the presence of binding motifs for TFs using HOMER (findMotifsGenome.pl) with RNAPII peaks outside the ecDNA targets as the background. Motifs results from HOMER were further selected based on the presence of the motifs in the background < 50% and *p*-value < 10^-12^. *Differential binding analysis*– DiffBind v3.8.4^70^ was used to perform differential MED1 binding. Briefly, consensus peaks generated from each of the samples were quantified using standard parameters after exclusion of MED1 peaks that matched chromosome M or Blacklist^71^. The DBA_DESEQ2 method was then applied to identify differential binding sites with the Benjamini– Hochberg false discovery rate (FDR) < 0.05^67^. For the comparisons of MED1 binding profiles, union of differential binding peaks (FDR ≤ 0.01) were used to calculate ChIP signals, ±1.2 kb or otherwise indicated, around MED1 peak summits with deeptools^72^. Heatmaps and averaged signal profiles were plotted using the deepTools functions plotHeatmap and plotProfile, respectively.

*Identification of SEs*– Super enhancers (SEs) and typical enhancers (TEs) were defined from the MED1 ChIP-seq peaks using the ROSE algorithm^32,73^ with the default parameters. Enhancers within a 12.5 kb genomic region were stitched to each other and ranked based on the density of ChIP-seq reads over input. SEs and TEs were classified according to the obtained plots of normalized MED1signals within enhancer regions versus enhancer ranks.

*Gene ontology (GO)* and *Gene set enrichment analysis (GSEA)*– GO enrichment analysis was performed by using either the PANTHER overrepresentation test (20240226 Release)^74^ or R package topGO (v2.52) with the “classic” algorithm. The *p*-values were determined using Fisher’s exact test with FDR correction. GSEA was conducted with 10,000 phenotype permutations using the function gseGO() [ont = “BP”] of R package ClusterProfiler (v4.8.3)^75^. All genes detected by RNA-seq were ranked according to log2FC from high to low. Gene sets with FDR < 0.05 were reported. Visualization of the GSEA results was performed using the R package “enrichplot”^76^.

### Imaging analyses

*Fluorescence in situ hybridization*– FISH analysis was performed as previously described^15^. In brief, metaphase cells were fixed in a methanol: acetic acid (3:1) solution and loaded on Poly-L-Lysine-coated slides (6776216, ThermoFisher Scientific). Cells were subsequently subjected to hybridization with the indicated FISH probes (MDM4-20-OR and EGFR-20-OR, Empire Genomics), washed, air dried, and stained with SlowFade Diamond Antifade Mounting medium with DAPI (S36964, ThermoFisher Scientific). Acquired images were processed using Fiji Is Just ImageJ (FIJI).

*Casilio live cell imaging and MED1 staining*– The CRISPR-based Casilio live cell imaging platform was performed as described^23^. Briefly, sgRNAs which have a built-in scaffold region containing 15 or 25 PUF domain binding sites (PBS) with specified 8-mer RNA sequences like PBSa and PBSc recruit programable RNA-binding proteins with PUF domains. The binding of multiple fluorescently labeled PUFs to the sgRNA scaffold enables an RNA-guided signal amplification at targeted sites. To image ecDNAs or chromosomal loci, cells were transfected with Casilio-imaging components containing 50 ng of dCas9 plasmid (Addgene #73169), 50 ng of Clover-PUFc plasmid (Addgene #73689), and 200 ng of the sgRNA-15xPBSc plasmid (derived from Addgene #183213) targeting the selected locus. In dual-label experiments, 300 ng of PUF9R-mRuby2 plasmid (Addgene #183210) and 200 ng of sgRNA-25xPBS9R plasmid (derived from Addgene #183217) were included with the Casilio-imaging components. Transfected cells were subjected to imaging 48 hours post-transfection, or as otherwise indicated, after staining nuclei with Hoechst 33342 (H3570, ThermoFisher Scientific).

Live imaging was performed at humidified 37°C 5% CO_2,_ on the Dragonfly High Speed Confocal Platform 505 (Andor) with a Leica DMi8 inverted microscope equipped with a live-cell environmental chamber (Okolab), an iXon EMCCD camera, and a Leica HC PL APO 63x/1.47NA OIL CORR TIRF objective. Images were acquired and processed as previously reported^23^.

For dual-label imaging of ecDNAs and their chromosomal targets, cells were transfected with Casilio-imaging components targeting ecDNA-interacting chromosomal loci or the non-interacting regions from the same chromosomes (controls). The distances between ecDNA spots and target genes as well as control loci were determined. The “density” function (in R version 4.2.1) was used to calculate the kernel density estimation of each given ecDNA-target or ecDNA-control pair to generate density distribution plots with an upper limit distance filter cutoff of 1.2 µm. Since the distribution is non-normal, we used a non-parametric statistical test, i.e. Wilcoxon rank sum test, to obtain the significance if a higher density of ecDNAs surrounding a target was observed versus that of the respective control locus.

To determine the sub-nuclear localization of ecDNA puncta and MED1, cells were subjected to immunofluorescence staining using MED1 antibody 48 hours after transfection with Casilio-imaging components. Cells were fixed, permeabilized, and treated with rabbit anti-MED1 primary antibody (ab64965, Abcam) and Alexa Fluor 568 conjugated goat anti-rabbit secondary antibody (A-11036, ThermoFisher Scientific). Hoechst 33342 was used to stain nuclei. Using Fiji image analysis software^77^, all pairwise 3D distances were measured from both Clover and Alexa Fluor 568 images by JACoP plugin^78^. For each nucleus, distances from each *ecMYC* spot to the nearest MED1 signals were averaged by performing objects -based methods using thresholds of 3,200 and 2,000 for Clover and Alexa Fluor 568 images, respectively. To evaluate if the distributions of the observed distances between *ecMYC* and MED1 foci were significantly shorter than random in a confined system with simulating uniform distributions, we fitted the nucleus to an ellipsoid using 8-bit Hoechst images and the 3D Fiji Suite plugin^79^. Fitted ellipsoid vectors, 10% extended radii, were used to generate 100,000 random points with uniform distribution. For each random point, the closest distance to MED1 was calculated. Constrained by collision, the closest distances of random points to MED1 < 0.2 µm were filtered out.

To image cells with 3% 1,6-HD treatment, cells were imaged under time-lapse microscopy immediately after the drug was added to the wells. Z-series at 0.4 µm step size and spanning the full nucleus was acquired every five or ten minutes. Dual-label imaging was performed using cells transfected with Casilio-imaging components containing sgRNAs targeting an unamplified control region. Plasmids with Addgene #73169, #73689, #183209, #183210, #183213, #183214, and # 183217 were gifts from Dr. Albert W. Cheng^23,42^.

For imaging ecDNA condensates in cells subjected to Casilio-i perturbations, Casilio-i cell lines were transfected with Casilio-imaging components, and images of ec*MYC* puncta were taken on the third day post Dox induction. Background fluorescence from Clover was used to normalize for transfection and the indicated nuclei counts were used to determine the fractions of nuclei containing ecDNA spots for each condition.

### Cell culture and functional characterization

K562 cells were grown in Iscoves medium supplemented with L-glutamine (12440053, ThermoFisher Scientific), 10% fetal bovine serum (FBS) (26140079, ThermoFisher Scientific), and 0.1 mM Non-Essential Amino Acids (11140050, ThermoFisher Scientific). PC3 DM and PC3 DM (-) cells were cultivated in F-12K (Kaighn’s modification of Ham’s F-12 medium) (30-2004, ATCC) supplemented with 10% FBS. COLO320 DM was grown in RPMI-1640 medium (30-2001, ATCC) supplemented with 10% FBS. GBM171 DM and GBM172 were cultivated in ultra-low attachment dishes (Corning) using DMEM/ Ham’s F-12 with GlutaMAX (10565018, ThermoFisher Scientific) supplemented with B27 supplement without vitamin A (12587010, ThermoFisher Scientific), 20 ng/ml FGF-Basic (154 a. a.) recombinant Human protein (100-18B, PeproTech), 20 ng/ml EGF recombinant Human protein (AF-100-15, Pepro Tech), 2.5 µg/ml heparin sodium salt (H3149-50KU, Sigma-Aldrich), and penicillin-streptomycin (15140122, ThermoFisher Scientific). All cell lines were tested for mycoplasma using MycoAlert Mycoplasma Detection Kit (LT07-418, Lonza).

*Drug treatment*– For each experiment, a 30% w/v stock solution of 1,6-HD (240117, Sigma-Aldrich,) was prepared in base media without serum. Cells were incubated for the indicated duration in the cell culture medium containing 3% 1,6-HD and were processed for subsequent analyses as described. For imaging analyses, 3% 1,6-HD was added directly to the wells of optical plates and images were acquired as described.

*Apoptosis assay*– Cells plated in a 96-well plate overnight were incubated for 30 min in the presence of CellEvent Caspase-3/7 Green reagent (R37111, Invitrogen). For quantification, cells were detached and moved to a V-bottom 96-well plate for flow cytometry analysis using a FACSymphony A5 flow cytometer (BD Bioscience) equipped with a high-throughput plate sampler and FACSDiVA software (BD Bioscience). Ten thousand cell events were collected for each sample to obtain average fluorescence intensities. Positive control of cells treated with 2 µg/mL actinomycin D for 24 hours were run in parallel for each experiment.

*Cell proliferation assay*– Cells were seeded in two sets of triplicates on a 96-well plate at 3,000 cells/well. 10 µL of CCK-8 (Cell Counting Kit-8) (96992, Sigma-Aldrich) was added in each well, and absorbance at 450 nm was measured as indicated by the manufacturer’s instructions using a SpectraMax M3 spectrophotometer (Molecular devices). Tetrazolium salts in CCK8, when mixed with cells, are reduced by cellular dehydrogenases to formazan products that can be measured by OD_abs_ 450 nm and whose cumulative amounts are directly proportional to the number of viable cells.

### CRISPR-based perturbation

*sgRNA design*– Due to the large genomic span of ecSEs, seven sgRNAs were designed per ecSE locus centered around the peaks of MED1-binding sites defined by ChIP-seq analysis. Their coordinates can be found in Supplementary Table 3. These sgRNAs were Golden Gate-assembled into a U6 promoter-based polycistronic plasmid to allow their co-transfection.

*Transfection*– K562-based CRISPR cell lines were nucleofected with 500 ng sgRNA plasmid DNA using 4-D nucleofector X Unit with 16-well nucleocuvette strips according to the manufacturer’s instructions (Lonza). For Casilio-i perturbations, cells were seeded in 24-well plates then transfected with 700 ng sgRNA or control plasmid DNAs using lipofectamine LTX and lipofectamine 3000 for PC3 DM and COLO320 DM cell lines, respectively, following the manufacturer’s instructions (15338030 and L3000008, ThermoFisher Scientific).

*Establishment of CRISPR stable cell lines*– Stable and Dox-inducible expression cell lines were generated by transfecting the indicated PiggyBac vectors in the presence of hyperactive transposase plasmid (hyPBase) as previously reported^80^. Transfected cells were subjected to four rounds of selection in the presence of blasticidin for CRISPRi or blasticidin and hygromycin for Casilio-based and enCRISPRi cells lines. Selected cells were further subjected to bulk cell sorting using a BD FACSAria cell sorter (BD Bioscience) according to the fluorescent label(s) used for the constructs to generate each of the cell lines. Plasmids with Addgene #138462 and 138460 were gifts from Dr. Jian Xu^41^ and used to generate the enCRISPRi used in this study. The expression of CRISPR-interference proteins in the established cell lines was further validated by Western blot analysis using anti-Cas9 (A-9000-100, Epigentek), anti-HA (H3663, Sigma-Aldrich), or anti-Flag (F1804, Sigma-Aldrich) antibodies and Clarity Western ECL Substrate (1705060, Bio-Rad). Dox-inducible Casilio-i stable cell lines derived from PC3 DM cells transiently transfected with sgRNA-plasmids targeting ecSE_1_, ecSE_combo_, or empty vector in the presence of Dox were subjected to puromycin selection for two days and then collected on day three post-transfection.

## References

1. Kim, H., Nguyen, N.P., Turner, K., Wu, S., Gujar, A.D., Luebeck, J., Liu, J., Deshpande, V., Rajkumar, U., Namburi, S., et al. (2020). Extrachromosomal DNA is associated with oncogene amplification and poor outcome across multiple cancers. Nat Genet 52, 891–897. 10.1038/s41588-020-0678-2.

2. Zhu, Y., Gong, L., and Wei, C.L. (2022). Guilt by association: EcDNA as a mobile transactivator in cancer. Trends Cancer 8, 747–758. 10.1016/j.trecan.2022.04.011.

3. deCarvalho, A.C., Kim, H., Poisson, L.M., Winn, M.E., Mueller, C., Cherba, D., Koeman, J., Seth, S., Protopopov, A., Felicella, M., et al. (2018). Discordant inheritance of chromosomal and extrachromosomal DNA elements contributes to dynamic disease evolution in glioblastoma. Nat Genet 50, 708–717. 10.1038/s41588-018-0105-0.

4. Verhaak, R.G.W., Bafna, V., and Mischel, P.S. (2019). Extrachromosomal oncogene amplification in tumour pathogenesis and evolution. Nat Rev Cancer 19, 283–288. 10.1038/s41568-019-0128-6.

5. Yan, X., Mischel, P., and Chang, H. (2024). Extrachromosomal DNA in cancer. Nat Rev Cancer 24, 261–273. 10.1038/s41568-024-00669-8.

6. Turner, K.M., Deshpande, V., Beyter, D., Koga, T., Rusert, J., Lee, C., Li, B., Arden, K., Ren, B., Nathanson, D.A., et al. (2017). Extrachromosomal oncogene amplification drives tumour evolution and genetic heterogeneity. Nature 543, 122–125. 10.1038/nature21356.

7. Nathanson, D.A., Gini, B., Mottahedeh, J., Visnyei, K., Koga, T., Gomez, G., Eskin, A., Hwang, K., Wang, J., Masui, K., et al. (2014). Targeted therapy resistance mediated by dynamic regulation of extrachromosomal mutant EGFR DNA. Science 343, 72–76. 10.1126/science.1241328.

8. Xu, K., Ding, L., Chang, T.C., Shao, Y., Chiang, J., Mulder, H., Wang, S., Shaw, T.I., Wen, J., Hover, L., et al. (2019). Structure and evolution of double minutes in diagnosis and relapse brain tumors. Acta Neuropathol 137, 123–137. 10.1007/s00401-018-1912-1.

9. Zheng, S., Fu, J., Vegesna, R., Mao, Y., Heathcock, L.E., Torres-Garcia, W., Ezhilarasan, R., Wang, S., McKenna, A., Chin, L., et al. (2013). A survey of intragenic breakpoints in glioblastoma identifies a distinct subset associated with poor survival. Genes Dev 27, 1462–1472. 10.1101/gad.213686.113.

10. Xue, Y., Martelotto, L., Baslan, T., Vides, A., Solomon, M., Mai, T.T., Chaudhary, N., Riely, G.J., Li, B.T., Scott, K., et al. (2017). An approach to suppress the evolution of resistance in BRAF(V600E)-mutant cancer. Nat Med 23, 929–937. 10.1038/nm.4369.

11. Yi, E., Gujar, A.D., Guthrie, M., Kim, H., Zhao, D., Johnson, K.C., Amin, S.B., Costa, M.L., Yu, Q., Das, S., et al. (2022). Live-Cell Imaging Shows Uneven Segregation of Extrachromosomal DNA Elements and Transcriptionally Active Extrachromosomal DNA Hubs in Cancer. Cancer Discov 12, 468–483. 10.1158/2159-8290.CD-21-1376.

12. Yi, E., Chamorro Gonzalez, R., Henssen, A.G., and Verhaak, R.G.W. (2022). Extrachromosomal DNA amplifications in cancer. Nat Rev Genet 23, 760–771. 10.1038/s41576-022-00521-5.

13. Wu, S., Turner, K.M., Nguyen, N., Raviram, R., Erb, M., Santini, J., Luebeck, J., Rajkumar, U., Diao, Y., Li, B., et al. (2019). Circular ecDNA promotes accessible chromatin and high oncogene expression. Nature 575, 699–703. 10.1038/s41586-019-1763-5.

14. Morton, A.R., Dogan-Artun, N., Faber, Z.J., MacLeod, G., Bartels, C.F., Piazza, M.S., Allan, K.C., Mack, S.C., Wang, X., Gimple, R.C., et al. (2019). Functional Enhancers Shape Extrachromosomal Oncogene Amplifications. Cell 179, 1330–1341 e1313. 10.1016/j.cell.2019.10.039.

15. Zhu, Y., Gujar, A.D., Wong, C.H., Tjong, H., Ngan, C.Y., Gong, L., Chen, Y.A., Kim, H., Liu, J., Li, M., et al. (2021). Oncogenic extrachromosomal DNA functions as mobile enhancers to globally amplify chromosomal transcription. Cancer Cell 39, 694–707 e697. 10.1016/j.ccell.2021.03.006.

16. Hung, K.L., Yost, K.E., Xie, L., Shi, Q., Helmsauer, K., Luebeck, J., Schopflin, R., Lange, J.T., Chamorro Gonzalez, R., Weiser, N.E., et al. (2021). ecDNA hubs drive cooperative intermolecular oncogene expression. Nature 600, 731–736. 10.1038/s41586-021-04116-8.

17. Hirose, T., Ninomiya, K., Nakagawa, S., and Yamazaki, T. (2023). A guide to membraneless organelles and their various roles in gene regulation. Nat Rev Mol Cell Biol 24, 288–304. 10.1038/s41580-022-00558-8.

18. Misteli, T. (2020). The Self-Organizing Genome: Principles of Genome Architecture and Function. Cell 183, 28–45. 10.1016/j.cell.2020.09.014.

19. Zheng, M., Tian, S.Z., Capurso, D., Kim, M., Maurya, R., Lee, B., Piecuch, E., Gong, L., Zhu, J.J., Li, Z., et al. (2019). Multiplex chromatin interactions with single-molecule precision. Nature 566, 558–562. 10.1038/s41586-019-0949-1.

20. Dixon, J.R., Selvaraj, S., Yue, F., Kim, A., Li, Y., Shen, Y., Hu, M., Liu, J.S., and Ren, B. (2012). Topological domains in mammalian genomes identified by analysis of chromatin interactions. Nature 485, 376–380. 10.1038/nature11082.

21. Mumbach, M.R., Rubin, A.J., Flynn, R.A., Dai, C., Khavari, P.A., Greenleaf, W.J., and Chang, H.Y. (2016). HiChIP: efficient and sensitive analysis of protein-directed genome architecture. Nat Methods 13, 919–922. 10.1038/nmeth.3999.

22. Quinn, L.A., Moore, G.E., Morgan, R.T., and Woods, L.K. (1979). Cell lines from human colon carcinoma with unusual cell products, double minutes, and homogeneously staining regions. Cancer Res 39, 4914–4924.

23. Clow, P.A., Du, M., Jillette, N., Taghbalout, A., Zhu, J.J., and Cheng, A.W. (2022). CRISPR-mediated multiplexed live cell imaging of nonrepetitive genomic loci with one guide RNA per locus. Nat Commun 13, 1871. 10.1038/s41467-022-29343-z.

24. Sondka, Z., Dhir, N.B., Carvalho-Silva, D., Jupe, S., Madhumita, McLaren, K., Starkey, M., Ward, S., Wilding, J., Ahmed, M., et al. (2024). COSMIC: a curated database of somatic variants and clinical data for cancer. Nucleic Acids Res 52, D1210–D1217. 10.1093/nar/gkad986.

25. Dressler, L., Bortolomeazzi, M., Keddar, M.R., Misetic, H., Sartini, G., Acha-Sagredo, A., Montorsi, L., Wijewardhane, N., Repana, D., Nulsen, J., et al. (2022). Comparative assessment of genes driving cancer and somatic evolution in non-cancer tissues: an update of the Network of Cancer Genes (NCG) resource. Genome Biol 23, 35. 10.1186/s13059-022-02607-z.

26. Kivioja, T., Vaharautio, A., Karlsson, K., Bonke, M., Enge, M., Linnarsson, S., and Taipale, J. (2011). Counting absolute numbers of molecules using unique molecular identifiers. Nat Methods 9, 72–74. 10.1038/nmeth.1778.

27. Sabari, B.R., Dall’Agnese, A., Boija, A., Klein, I.A., Coffey, E.L., Shrinivas, K., Abraham, B.J., Hannett, N.M., Zamudio, A.V., Manteiga, J.C., et al. (2018). Coactivator condensation at super-enhancers links phase separation and gene control. Science 361. 10.1126/science.aar3958.

28. Heinz, S., Benner, C., Spann, N., Bertolino, E., Lin, Y.C., Laslo, P., Cheng, J.X., Murre, C., Singh, H., and Glass, C.K. (2010). Simple combinations of lineage-determining transcription factors prime cis-regulatory elements required for macrophage and B cell identities. Mol Cell 38, 576–589. 10.1016/j.molcel.2010.05.004.

29. Yan, G., and Lei, W. (2023). Role of ELK1 in regulating colorectal cancer progression: miR-31-5p/CDIP1 axis in CRC pathogenesis. PeerJ 11, e15602. 10.7717/peerj.15602.

30. Liu, X., Chen, J., Zhang, S., Liu, X., Long, X., Lan, J., Zhou, M., Zheng, L., and Zhou, J. (2022). LINC00839 promotes colorectal cancer progression by recruiting RUVBL1/Tip60 complexes to activate NRF1. EMBO Rep 23, e54128. 10.15252/embr.202154128.

31. Takayama, K.I., Kosaka, T., Suzuki, T., Hongo, H., Oya, M., Fujimura, T., Suzuki, Y., and Inoue, S. (2021). Subtype-specific collaborative transcription factor networks are promoted by OCT4 in the progression of prostate cancer. Nat Commun 12, 3766. 10.1038/s41467-021-23974-4.

32. Whyte, W.A., Orlando, D.A., Hnisz, D., Abraham, B.J., Lin, C.Y., Kagey, M.H., Rahl, P.B., Lee, T.I., and Young, R.A. (2013). Master transcription factors and mediator establish super-enhancers at key cell identity genes. Cell 153, 307–319. 10.1016/j.cell.2013.03.035.

33. Geiger, F., Acker, J., Papa, G., Wang, X., Arter, W.E., Saar, K.L., Erkamp, N.A., Qi, R., Bravo, J.P., Strauss, S., et al. (2021). Liquid-liquid phase separation underpins the formation of replication factories in rotaviruses. EMBO J 40, e107711. 10.15252/embj.2021107711.

34. Zhang, Y., Wong, C.H., Birnbaum, R.Y., Li, G., Favaro, R., Ngan, C.Y., Lim, J., Tai, E., Poh, H.M., Wong, E., et al. (2013). Chromatin connectivity maps reveal dynamic promoter-enhancer long-range associations. Nature 504, 306–310. 10.1038/nature12716.

35. Yang, T., Zhang, F., Yardimci, G.G., Song, F., Hardison, R.C., Noble, W.S., Yue, F., and Li, Q. (2017). HiCRep: assessing the reproducibility of Hi-C data using a stratum-adjusted correlation coefficient. Genome Res 27, 1939–1949. 10.1101/gr.220640.117.

36. Maity, G., Mehta, S., Haque, I., Dhar, K., Sarkar, S., Banerjee, S.K., and Banerjee, S. (2014). Pancreatic tumor cell secreted CCN1/Cyr61 promotes endothelial cell migration and aberrant neovascularization. Sci Rep 4, 4995. 10.1038/srep04995.

37. Riya, P.A., Basu, B., Surya, S., Parvathy, S., Lalitha, S., Jyothi, N.P., Meera, V., Jaikumar, V.S., Sunitha, P., Shahina, A., et al. (2022). HES1 promoter activation dynamics reveal the plasticity, stemness and heterogeneity in neuroblastoma cancer stem cells. J Cell Sci 135. 10.1242/jcs.260157.

38. Seim, I., Jeffery, P.L., Thomas, P.B., Nelson, C.C., and Chopin, L.K. (2017). Whole-Genome Sequence of the Metastatic PC3 and LNCaP Human Prostate Cancer Cell Lines. G3 (Bethesda) 7, 1731-1741. 10.1534/g3.117.039909.

39. Ueda, J., Yamazaki, T., and Funakoshi, H. (2023). Toward the Development of Epigenome Editing-Based Therapeutics: Potentials and Challenges. Int J Mol Sci 24. 10.3390/ijms24054778.

40. Fulco, C.P., Munschauer, M., Anyoha, R., Munson, G., Grossman, S.R., Perez, E.M., Kane, M., Cleary, B., Lander, E.S., and Engreitz, J.M. (2016). Systematic mapping of functional enhancer-promoter connections with CRISPR interference. Science 354, 769–773. 10.1126/science.aag2445.

41. Li, K., Liu, Y., Cao, H., Zhang, Y., Gu, Z., Liu, X., Yu, A., Kaphle, P., Dickerson, K.E., Ni, M., and Xu, J. (2020). Interrogation of enhancer function by enhancer-targeting CRISPR epigenetic editing. Nat Commun 11, 485. 10.1038/s41467-020-14362-5.

42. Cheng, A.W., Jillette, N., Lee, P., Plaskon, D., Fujiwara, Y., Wang, W., Taghbalout, A., and Wang, H. (2016). Casilio: a versatile CRISPR-Cas9-Pumilio hybrid for gene regulation and genomic labeling. Cell Res 26, 254–257. 10.1038/cr.2016.3.

43. Nunez, J.K., Chen, J., Pommier, G.C., Cogan, J.Z., Replogle, J.M., Adriaens, C., Ramadoss, G.N., Shi, Q., Hung, K.L., Samelson, A.J., et al. (2021). Genome-wide programmable transcriptional memory by CRISPR-based epigenome editing. Cell 184, 2503–2519 e2517. 10.1016/j.cell.2021.03.025.

44. Yeo, N.C., Chavez, A., Lance-Byrne, A., Chan, Y., Menn, D., Milanova, D., Kuo, C.C., Guo, X., Sharma, S., Tung, A., et al. (2018). An enhanced CRISPR repressor for targeted mammalian gene regulation. Nat Methods 15, 611–616. 10.1038/s41592-018-0048-5.

45. Kaya-Okur, H.S., Wu, S.J., Codomo, C.A., Pledger, E.S., Bryson, T.D., Henikoff, J.G., Ahmad, K., and Henikoff, S. (2019). CUT&Tag for efficient epigenomic profiling of small samples and single cells. Nat Commun 10, 1930. 10.1038/s41467-019-09982-5.

46. Cho, S.W., Xu, J., Sun, R., Mumbach, M.R., Carter, A.C., Chen, Y.G., Yost, K.E., Kim, J., He, J., Nevins, S.A., et al. (2018). Promoter of lncRNA Gene PVT1 Is a Tumor-Suppressor DNA Boundary Element. Cell 173, 1398–1412 e1322. 10.1016/j.cell.2018.03.068.

47. Chai, H., Tjong, H., Li, P., Liao, W., Wang, P., Wong, C.H., Ngan, C.Y., Leonard, W.J., Wei, C.L., and Ruan, Y. (2023). ChIATAC is an efficient strategy for multi-omics mapping of 3D epigenomes from low-cell inputs. Nat Commun 14, 213. 10.1038/s41467-023-35879-5.

48. Yaqinuddin, A., Qureshi, S.A., Qazi, R., and Abbas, F. (2008). Down-regulation of DNMT3b in PC3 cells effects locus-specific DNA methylation, and represses cellular growth and migration. Cancer Cell Int 8, 13. 10.1186/1475-2867-8-13.

49. Zhu, A., Hopkins, K.M., Friedman, R.A., Bernstock, J.D., Broustas, C.G., and Lieberman, H.B. (2021). DNMT1 and DNMT3B regulate tumorigenicity of human prostate cancer cells by controlling RAD9 expression through targeted methylation. Carcinogenesis 42, 220–231. 10.1093/carcin/bgaa088.

50. Ji, J., Shen, T., Li, Y., Liu, Y., Shang, Z., and Niu, Y. (2021). CDCA5 promotes the progression of prostate cancer by affecting the ERK signalling pathway. Oncol Rep 45, 921–932. 10.3892/or.2021.7920.

51. Miyamoto, T., Min, W., and Lillehoj, H.S. (2002). Lymphocyte proliferation response during Eimeria tenella infection assessed by a new, reliable, nonradioactive colorimetric assay. Avian Dis 46, 10–16. 10.1637/0005-2086(2002)046[0010:LPRDET]2.0.CO;2.

52. Shin, Y., and Brangwynne, C.P. (2017). Liquid phase condensation in cell physiology and disease. Science 357. 10.1126/science.aaf4382.

53. Cremer, T., and Cremer, M. (2010). Chromosome territories. Cold Spring Harb Perspect Biol 2, a003889. 10.1101/cshperspect.a003889.

54. Maass, P.G., Barutcu, A.R., and Rinn, J.L. (2019). Interchromosomal interactions: A genomic love story of kissing chromosomes. J Cell Biol 218, 27–38. 10.1083/jcb.201806052.

55. Bashkirova, E., and Lomvardas, S. (2019). Olfactory receptor genes make the case for inter-chromosomal interactions. Curr Opin Genet Dev 55, 106–113. 10.1016/j.gde.2019.07.004.

56. Monahan, K., Horta, A., and Lomvardas, S. (2019). LHX2- and LDB1-mediated trans interactions regulate olfactory receptor choice. Nature 565, 448–453. 10.1038/s41586-018-0845-0.

57. Moreau, P., Cournac, A., Palumbo, G.A., Marbouty, M., Mortaza, S., Thierry, A., Cairo, S., Lavigne, M., Koszul, R., and Neuveut, C. (2018). Tridimensional infiltration of DNA viruses into the host genome shows preferential contact with active chromatin. Nat Commun 9, 4268. 10.1038/s41467-018-06739-4.

58. Hensel, K.O., Cantner, F., Bangert, F., Wirth, S., and Postberg, J. (2018). Episomal HBV persistence within transcribed host nuclear chromatin compartments involves HBx. Epigenetics Chromatin 11, 34. 10.1186/s13072-018-0204-2.

59. Lozzio, B.B., and Lozzio, C.B. (1979). Properties and usefulness of the original K-562 human myelogenous leukemia cell line. Leuk Res 3, 363–370. 10.1016/0145-2126(79)90033-x.

60. Mao, D.D., Gujar, A.D., Mahlokozera, T., Chen, I., Pan, Y., Luo, J., Brost, T., Thompson, E.A., Turski, A., Leuthardt, E.C., et al. (2015). A CDC20-APC/SOX2 Signaling Axis Regulates Human Glioblastoma Stem-like Cells. Cell Rep 11, 1809–1821. 10.1016/j.celrep.2015.05.027.

61. Gujar, A.D., Mao, D.D., Finlay, J.B., and Kim, A.H. (2018). Establishing Primary Human Glioblastoma Adherent Cultures from Operative Specimens. Methods Mol Biol 1741, 53-62. 10.1007/978-1-4939-7659-1_3.

62. Miller, C.A., Hampton, O., Coarfa, C., and Milosavljevic, A. (2011). ReadDepth: a parallel R package for detecting copy number alterations from short sequencing reads. PLoS One 6, e16327. 10.1371/journal.pone.0016327.

63. Wang, P., Feng, Y., Zhu, K., Chai, H., Chang, Y.T., Yang, X., Liu, X., Shen, C., Gega, E., Lee, B., et al. (2021). In situ Chromatin Interaction Analysis Using Paired-End Tag Sequencing. Curr Protoc 1, e174. 10.1002/cpz1.174.

64. Paulsen, J., Rodland, E.A., Holden, L., Holden, M., and Hovig, E. (2014). A statistical model of ChIA-PET data for accurate detection of chromatin 3D interactions. Nucleic Acids Res 42, e143. 10.1093/nar/gku738.

65. Kaufmann, S., Fuchs, C., Gonik, M., Khrameeva, E.E., Mironov, A.A., and Frishman, D. (2015). Inter-chromosomal contact networks provide insights into Mammalian chromatin organization. PLoS One 10, e0126125. 10.1371/journal.pone.0126125.

66. Kruse, K., Sewitz, S., and Babu, M.M. (2013). A complex network framework for unbiased statistical analyses of DNA-DNA contact maps. Nucleic Acids Res 41, 701–710. 10.1093/nar/gks1096.

67. Love, M.I., Huber, W., and Anders, S. (2014). Moderated estimation of fold change and dispersion for RNA-seq data with DESeq2. Genome Biol 15, 550. 10.1186/s13059-014-0550-8.

68. Kaya-Okur, H.S., Janssens, D.H., Henikoff, J.G., Ahmad, K., and Henikoff, S. (2020). Efficient low-cost chromatin profiling with CUT&Tag. Nat Protoc 15, 3264–3283. 10.1038/s41596-020-0373-x.

69. Kim, D., Langmead, B., and Salzberg, S.L. (2015). HISAT: a fast spliced aligner with low memory requirements. Nat Methods 12, 357–360. 10.1038/nmeth.3317.

70. Ross-Innes, C.S., Stark, R., Teschendorff, A.E., Holmes, K.A., Ali, H.R., Dunning, M.J., Brown, G.D., Gojis, O., Ellis, I.O., Green, A.R., et al. (2012). Differential oestrogen receptor binding is associated with clinical outcome in breast cancer. Nature 481, 389–393. 10.1038/nature10730.

71. Amemiya, H.M., Kundaje, A., and Boyle, A.P. (2019). The ENCODE Blacklist: Identification of Problematic Regions of the Genome. Sci Rep 9, 9354. 10.1038/s41598-019-45839-z.

72. Ramirez, F., Dundar, F., Diehl, S., Gruning, B.A., and Manke, T. (2014). deepTools: a flexible platform for exploring deep-sequencing data. Nucleic Acids Res 42, W187–191. 10.1093/nar/gku365.

73. Loven, J., Hoke, H.A., Lin, C.Y., Lau, A., Orlando, D.A., Vakoc, C.R., Bradner, J.E., Lee, T.I., and Young, R.A. (2013). Selective inhibition of tumor oncogenes by disruption of super-enhancers. Cell 153, 320–334. 10.1016/j.cell.2013.03.036.

74. Mi, H., Muruganujan, A., Ebert, D., Huang, X., and Thomas, P.D. (2019). PANTHER version 14: more genomes, a new PANTHER GO-slim and improvements in enrichment analysis tools. Nucleic Acids Res 47, D419–D426. 10.1093/nar/gky1038.

75. Subramanian, A., Tamayo, P., Mootha, V.K., Mukherjee, S., Ebert, B.L., Gillette, M.A., Paulovich, A., Pomeroy, S.L., Golub, T.R., Lander, E.S., and Mesirov, J.P. (2005). Gene set enrichment analysis: a knowledge-based approach for interpreting genome-wide expression profiles. Proc Natl Acad Sci U S A 102, 15545–15550. 10.1073/pnas.0506580102.

76. Wu, T., Hu, E., Xu, S., Chen, M., Guo, P., Dai, Z., Feng, T., Zhou, L., Tang, W., Zhan, L., et al. (2021). clusterProfiler 4.0: A universal enrichment tool for interpreting omics data. Innovation (Camb) 2, 100141. 10.1016/j.xinn.2021.100141.

77. Schindelin, J., Arganda-Carreras, I., Frise, E., Kaynig, V., Longair, M., Pietzsch, T., Preibisch, S., Rueden, C., Saalfeld, S., Schmid, B., et al. (2012). Fiji: an open-source platform for biological-image analysis. Nat Methods *9*, 676-682. 10.1038/nmeth.2019.

78. Bolte, S., and Cordelieres, F.P. (2006). A guided tour into subcellular colocalization analysis in light microscopy. J Microsc 224, 213–232. 10.1111/j.1365-2818.2006.01706.x.

79. Ollion, J., Cochennec, J., Loll, F., Escude, C., and Boudier, T. (2013). TANGO: a generic tool for high-throughput 3D image analysis for studying nuclear organization. Bioinformatics 29, 1840–1841. 10.1093/bioinformatics/btt276.

80. Yusa, K., Zhou, L., Li, M.A., Bradley, A., and Craig, N.L. (2011). A hyperactive piggyBac transposase for mammalian applications. Proc Natl Acad Sci U S A 108, 1531–1536. 10.1073/pnas.1008322108.

